# Single-cell transcriptomic landscape deciphers novel olfactory neuroblastoma subtypes and intratumoral heterogeneity

**DOI:** 10.1101/2023.01.24.522680

**Authors:** Jingyi Yang, Xiaole Song, Huankang Zhang, Quan Liu, Ruoyan Wei, Luo Guo, Cuncun Yuan, Kai Xue, Yuting Lai, Li Wang, Li Hu, Huan Wang, Chen Zhang, Qianqian Zhang, Ye Gu, Weidong Zhao, Huapeng Yu, Jingjing Wang, Zhuofu Liu, Han Li, Shixing Zheng, Juan Liu, Lu Yang, Wanpeng Li, Rui Xu, Jiani Chen, Yumin Zhou, Xiankui Cheng, Yiqun Yu, Dehui Wang, Xicai Sun, Hongmeng Yu

## Abstract

Olfactory neuroblastoma (ONB) is an uncommon malignant tumor known to originate from the olfactory epithelial. The complex tumor ecosystem of this pathology remained unclear. Here, we explored the cellular components with ONB tumors based on scRNA profiles of 96,325 single-cells derived from 10 tumors and 1 olfactory mucosa sample. We discovered 11 major cell types, including 6 immune cell, 3 stromal cell subtypes and epithelial cluster in the tumor microenvironment (TME), and identified 5 common expression programs from malignant epithelial cells. We analyzed subclusters of TME and the interactions among different cell types in the TME. An innovative three-classification of ONB was established via scRNA analysis. Markers for categorizing tumor samples into new subtypes were elucidated. Different responses towards certain chemotherapy regimens could be inferred according to the molecular features of three tumor types. Relative abundance of immunosuppressive TAMs indicated the benefits of immunotherapies targeting myeloid cells.

## Introduction

Olfactory neuroblastoma (ONB) is an uncommon malignant tumor that originates in the nasal vault^1^, which comprises only 5%-6% among all sinonasal malignant neoplasms^2^ with a sparse incidence of 0.4 cases per million/year^3^. Also known as esthesioneuroblastoma (ENB), the varying names for this rare malignancy is essentially attributed to its uncertain histological origin. Although recently ONB was increasingly proposed to arise from the olfactory mucosa in the nasal cleft^4^ the precise cellular component of olfactory epithelial from which ONB is generated remains await to be elucidated.

Despite the rarity, varying biological activity of ONB has been reported, ranging from indolent growth to a highly aggressive progression^1^. Orbital, dural and intracranial invasion were frequently observed since the neoplasm is often found to infract the anterior skull base^5^. Multi-modality management has been recommended for ONB patients, with radical resection followed by postoperative radiotherapy as the most promoted treatment strategy for primary cases^6^. The role and standard regimes of chemotherapy for ONB patients were yet to be ascertained. While the 10-year overall survival rate and 5-year disease free survival rate of ONB patients are approximately 50% and 40%^1, 7^, a review of literature revealed limited research findings from oncogenesis, molecular features to tumor microenvironment (TME) components of this uncommon pathology^4^.

The rapid development of high-throughput single-cell RNA sequencing (scRNA-seq) has provided valid methods to dissect both the atlas of diverse TME cell types and the heterogeneity of tumor cells. Moreover, scRNA-seq technologies could further assist deciphering the transcriptome dynamic, biological function and cellular interactions between malignant cells and tumor-infiltrating TME components, from which latent oncological pathogenesis and novel targets for potential drugs could be possibly found^8^.

Although recently a few studies have attempted to develop molecular-based subtype classifications and analysis transcriptional pathways with prognostic values of ONB based on formalin-fixed paraffin-embedded (FFPE) tumor blocks or bulk tissue RNA-seq^9, 10^, the results failed to precisely decode the transcriptome features of this tumor, since bulk-RNA seq could only detect mixed bio-information contributed both by malignant and tumor-infiltrating TME cells. Moreover, the dynamic status and mutual interaction networks inside various cellular components also remained ill-defined.

Herein, in the present study, we performed scRNA-seq with ONB tumor samples derived from 10 treatment-naive patients, trying to provide the first cellular landscape illustrating both intra- and inter-tumoral heterogeneity for ONB. Moreover, combined analysis with scRNA data obtained from normal adult olfactory mucosa brought us an opportunity to take a glimpse at the possible oncogenic origin of this tumor. By integrating our findings, we identified a novel malignant cell subtype with distinct transcriptomic performance, as well as several drug-targeted functional pathways of ONB cellular components, which might provide new insights and hence possibly inspire some new diagnostic and therapeutic strategies of this rare, uncommon malignancy.

## Results

### Single-cell expression atlas and cell type annotation in ONB and normal adult OM

To decipher the cellular atlas within the ONB tumors, the single-cell RNA-seq (scRNA) profiles of 10 ONB samples obtained during endoscopic tumor resection at the Eye, Ear, Nose, Throat Hospital of Fudan University, as well as one adult human OM sample retrieved from GEO dataset (GSE139522)^11^, were enrolled in this study following the intuitional workflow **(Figure 1a, Table S1)**. The fresh biopsies were rapidly digested into single-cell suspensions after surgical resection and then processed for 3’ droplet-based single-cell RNA-seq (10× Genomics). After several quality control (QC) steps (see **Materials and Methods**), we obtained transcriptional profiles of N = 96325 cells in total, including N = 85452 from 10 ONB samples and N = 10873 cells from the control OM samples **(Figure S1a, Table S2).**

**Figure 1.**
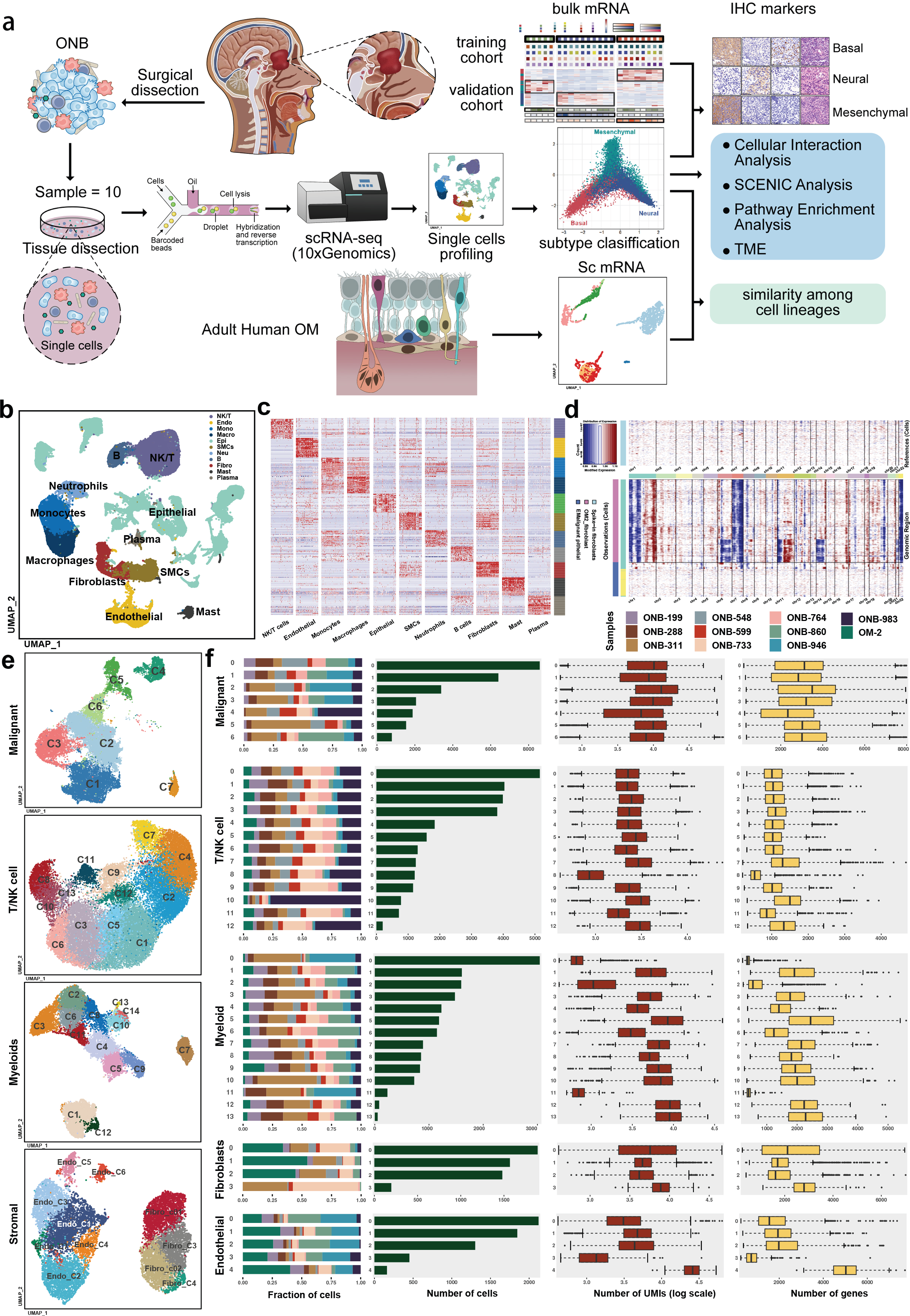
Single cell transcriptomic atlas of diverse cell types in ONB. **a)** A workflow depicting the overall experimental design for scRNA-seq profiling of ONB. **b)** UMAP visualization of 96325 single cells from 10 ONB samples and one normal adult olfactory mucosa sample (OM2), colored by different major cell types. **c)** Heatmap displaying expression levels of top 50 differentially expressed genes (DEGs) in each cell type displayed in **a**. Detailed gene list for DEGs were provided in **Supplementary Table S3**. **d)** Large-scale CNVs inferred based on scRNA-seq data of single cells from a representative tumor (ONB-599). Amplifications (red) or deletions (blue) were inferred by averaging expression over 100-gene stretches on the indicated chromosomes. Malignant epithelial cells were distinguished from non-malignant epithelial cells by distinctive patterns of CNV levels. **e)** UMAP plots showing the subclusters of malignant epithelial cells, T/NK cells, myeloid cells and stromal cells including ECs and fibroblasts. **f)** Detailed cellular component data of 7 malignant subgroups, 27 immune subclusters and 9 stromal subclusters from 11 samples (from left to right): the fraction of cells extracted from each sample, the number of cells, and box plots of the number of UMIs (log scale) and genes (with plot center, box and whiskers corresponding to median, IQR, 1.5 × IQR, and outliers, respectively).

Dimensional reduction was performed based on principle component analysis (PCA), followed by graph-based clustering, after which we identified 11 major cell clusters **(Figure 1b, Figure S1b)**. The cell subtypes were annotated according to canonical markers in the CellMarker database and olfactory neuronal markers previously described by Durante et al and Fletcher et al ^11–14^. **(Figure S1c)** Cell clusters were classified into 6 immune lineages (CD45^+^) including macrophages, dendric cells (DCs), monocytes, mast cells, T and B cells, 3 non-immune stromal lineages including fibroblasts, endothelial cells (ECs) and smooth muscle cells (SMCs) and one epithelial cluster marked by classic epithelial markers including EPCAM, CLDNs, and cytokeratin **(Figure 1c, Table S3)**. Fibroblasts (5529 cells, 5.7%) were marked with ACTA2 and COL1A2. The B cells (3338 cells, 3.5%) were marked with MS4A1 and CD79A, NK/T cells (26996 cells, 28.0%) were marked with CD2, CD3D and CD3E, macrophages (5723 cells, 5.9%) were marked with CD14 and the mast cells (1278 cells, 1.3%) were marked with CPA3.

Then we adopted a two-step approach based mainly on inferred CNV levels to identify malignant cells within the epithelial cluster **(Figure 1d)**^15, 16^. First, large-scale chromosomal copy number variations (CNVs) for each single-cell in the epithelial cluster were inferred according to averaged expression patterns across intervals of the genome. All the epithelial cells were selected as input and all the fibroblast identified in the OM dataset were extracted as control samples. Secondly, hierarchical clustering of the CNV profiles of the fibroblasts selected from the OM dataset and epithelial cells from each ONB tumor sample was performed separately (Figure S1d). Epithelial cells from the ONB dataset which inferred with polyploidic CNV patterns were identified as malignant cells, whereas cells which otherwise clustered together with spike-in fibroblasts from the normal OM were considered non-malignant. As a result, 24686 cells from 10 ONB tumor samples were designated as malignant cells and were extracted for further analyses.

Next, we performed extraction and re-clustering of each major cell type (malignant, myeloids, T/NK cells, fibroblasts and ECs) to identify subclusters. Attributes including cellular abundance, number of genes and UMIs, distribution of subgroups proportion in each patient, varied substantially among different tissue origins and patients **(Figure 1e-1f)**. The result revealed a highly heterogeneous ONB tumor environment (TME) in need of further illustration in the following analysis.

### Intra-tumoral Expression Heterogeneity of ONB malignant Cells

In order to depict the cellular diversity within the malignant compartment of ONB, tumoral epithelial cells were identified based on their elevated CNV levels compared to a subset of fibroblasts selected from normal olfactory mucosa as reference data (See **Material & Methods**; **Figure 1c**). As a result, a total of N = 24686 ONB cancer cells were screened out and re-clustered into 7 malignant subgroups **(Figure 2a)**. According to a multi-omics study of ONB^9^, two subtypes of this tumor, namely Basal and Neural, could be distinguished by unique molecular expression patterns and distinct clinicopathological features, with possibly diverse cell origins from different layers of olfactory epithelium^10^. To validated this binary classification of ONB at single-cell resolution, we firstly evaluated the expression levels of Basal/Neural signatures provided in the multi-omics research **(Table S4)**, with subtype-specific expression assessed by scores displayed in feature-plots **(Figure 2b)**. Preliminarily, Neural and Basal Scores preferentially distributed in different area of tumor UMAP, which demonstrated the existence of Basal and Neural-like malignant cells, while unexpectedly, we found that certain malignant subgroups (C4-C7) harbored a distinct expression pattern that could hardly be characterized using signatures of either known ONB subtypes **(Figure 2b, Figure S2a)**.

**Figure 2.**
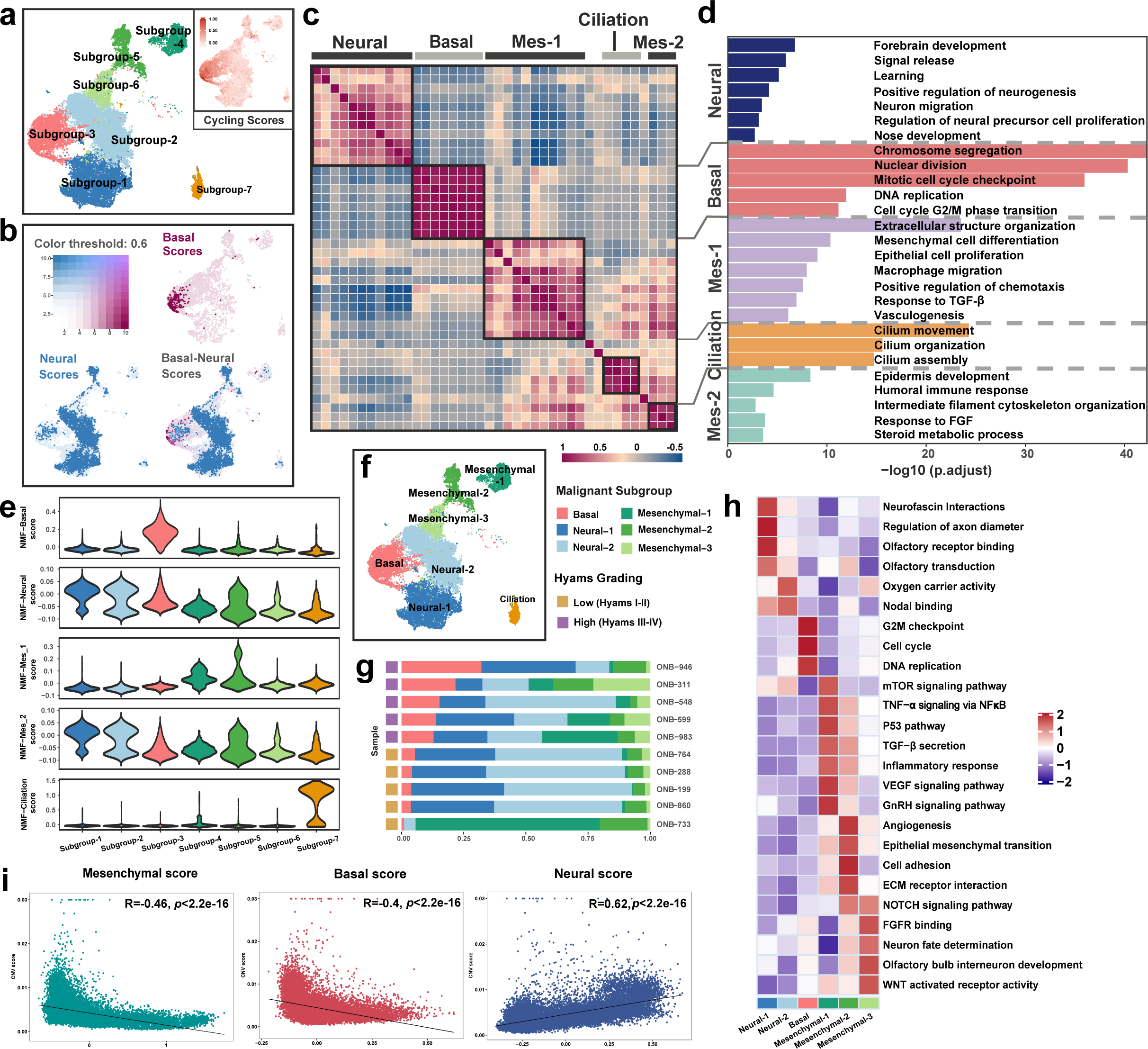
Intra-tumoral heterogeneity of ONB. **a)** UMAP representation of 7 subgroups generated from sub-clustering malignant epithelial cells. Cycling scores of each single cell were calculated using cell cycling-related gene list **(Table S4)**. **b)** UMAP depicting feature scores of canonical Basal and Neural ONB subtypes in malignant epithelial cells. **c)** Pairwise correlations of 40 intra-tumoral NMF programs derived from 10 ONB tumors as in **Table S5**. Hierarchical clustering identified 5 coherent meta-programs across tumors. **d)** Bar plots displaying the representative GO pathway terms enriched in each NMF meta-program. **e)** Panels of violin plots showing module scores for each of the 5 NMF meta-programs in 7 malignant subgroups (panels from top to bottom: Basal, Neural, Mes-1, Mes-2, Ciliation). **f)** UMAP displaying annotated phenotypes of 7 ONB malignant subgroups, noted on the color key legend and labels. **g)** Piled bar plots representing the cellular proportion of each malignant subgroup found in each ONB sample. **h)** Heatmap showing the representative GSVA pathway terms enriched in ONB malignant subgroups except for the Ciliation subcluster. Color key indicates z-score of -Log10(P value). **i)** The Pearson correlation analysis of inferred CNV levels (y-axis) and Neural, Basal, Mesenchymal signature scores in ONB malignant epithelial cells, respectively. The two-sided p value for each malignant signature was extremely small (p<2.22e×10^−15^), therefore the exact p values cannot be shown in the figure.

Non-negative matrix factorization (NMF) analysis was performed to investigate the expression features underlying the intra-tumoral heterogeneity of ONB (See **Material & Methods**; **Figure S2b, Table S5)**^17–19^. From each tumor dataset, 4 programs were extracted to generate a matrix of 40 intra-tumoral programs, which were distilled into 5 main meta-gene modules, after correlation hierarchical clustering **(Figure 2c, Table S6)**. The 5 meta-gene modules showed diverse cellular functions as annotated by gene ontology (GO) pathway analysis performed using top-scoring genes of each meta-program **(Figure 2d)**. Overall, we identified meta-programs enriched for olfactory neurodevelopment (meta-program 1; CHGB/HES6/NEUROD1) and cell cycling (meta-program 2; MKI67/TOP2A/UBE2C) pathways, which respectively reflected expression signatures of classical neural and basal ONB subtypes. Meta-program 4 (CAPS/TPPP3/PIFO) was found to associate with cilium movement and organization^20^. Moreover, functional analysis revealed relations between 2 NMF modules (meta-program 3, NRG1/WNT10A; meta-program 5, BMP7/LAMA1) and tumoral mesenchyme. The meta-programs were thus named Mes-1 and Mes-2, owing to their enrichment for pathways including extracellular structure organization, mesenchymal cell differentiation, responses to TGF-β and FGF, and metabolic-related processes. **(Table S7)**

The expression level of NMF meta-programs were evaluated in forms of module scores for each ONB malignant cell **(Figure 2e, Figure S2c)**. Overall, based on different expression patterns of these meta-programs, we were able to divide ONB malignant subgroups into 4 categories **(Figure 2f)**. The Neural and Basal meta-programs were found respectively enriched in subgroups C1-C2 and subgroup C3, which reflected the features of canonical Neural/Basal ONB subtypes. Subgroup C7 were identified as ciliated cancer cells due to distinctive expression of the Ciliation meta-program, as previously recognized in researches of scRNA profiling on recurrent nasopharyngeal carcinoma and lung cancers^20, 21^. Notably, malignant subgroups C4-C6 were regarded as mesenchymal-like subpopulations due to their relative enrichment for Mes-1 and Mes-2 meta-programs.

The identification of mesenchymal-like malignant subgroups sparked our interest in getting a further insight into intra-tumoral heterogeneity of ONB. The cellular proportion of Basal subgroup was found to be positively correlated with higher Hyams grades (Hyams III&IV, **Figures 2g**). GSVA enrichment analysis was performed to reveal the functional roles played by malignant subgroups **(Figure 2h)**. The Basal subgroup (C3) cells, enriched for proliferation pathways and high levels of Cell Cycle Scores **(Figure 2a)**, were regarded as the proliferative subpopulation in ONB tumor compartment. Pathways mainly related to neurogenesis and olfactory transduction were found up-regulated in the Neural subgroups (C1&C2). Specifically, the Mesenchymal subgroups (C4-C6) displayed multiple functions corresponding cellular activities in TME, ranging from angiogenesis, epithelial mesenchymal transition (EMT), extra cellular matrix (ECM) remolding, to mTOR signaling, inflammatory response and chemokine secretion, as well as olfactory bulb interneuron development, indicating a broad interaction of the Mesenchymal subpopulations with ONB ecosystems.

The differentially expressed genes (DEGs) among Neural/Basal/Mesenchymal (NBM) malignant cells were calculated, and the top 50 up-regulated DEGs were used to evaluate malignant scores in each ONB tumor cell **(Table S8)**. An uneven distribution pattern of inferred CNV scores across NBM subpopulations was discovered. For instance, we found a tendency of positive association between inferred CNVs and Neural Scores, and in opposite, a negative correlation with CNV levels shared by both Mesenchymal and Basal Scores in ONB malignant cells **(Figure 2i, Figure S2d-S2e)**. Considering a higher mutation rates of the basal subtype reported previously, our results indicated that the extent of putative copy number alterations was probably unparallel with mutational load when evaluated across different ONB subtypes^22^.

### Projection and Validation of Neural/Basal/Mesenchymal Signatures Stratifying Bulk ONB Samples

In order to further illustrate the spectrum reflecting hybrid cellular states within individual ONB samples at single-cell level, we created a ternary diagram, with malignant cells except the cilia subgroup (C7) placed on the plot according to their relative scores regarding NBM classifier genes, with different categories (Neural/Basal/Mesenchymal subpopulations) marked by distinct colors (see **Material & Methods, Figure 3a)**. Overall, our ternary diagram confirmed a tripartite structure of cellular phenotypes across ONB malignant cells. However, when splitting the ternary map to display NBM cellular compositions of each ONB patient, we discovered that some tumors favored particular cellular states over other categories, demonstrating 3 distinguishable patterns **(Figure 3b)**. For instance, in 5 poorly differentiated tumors with high Hyams grades (III-IV), the Basal malignant cells occupied a main status in tumor compartment, accompanied by a Mesenchymal subpopulation and a group of low-scoring Neural cells. Furthermore, we identified four cases with relatively low Hyams grads (I-II) which consisted primarily of high-scoring Neural cells and a subset of moderately-scored Mesenchymal cell. These ONB samples, dominated by Basal and Neural subpopulations respectively, were considered equivalent to the canonical Basal and neural ONB subtypes^9^. Moreover, we noticed that there was one tumor sample (ONB-733) comprised of almost only Mesenchymal cells, implying a possible existence of a novel ONB subtype featured by exclusive expression of Mesenchymal signatures, whereas holding inadequate proof of either Basal or Neural-like expression patterns in the meanwhile.

**Figure 3.**
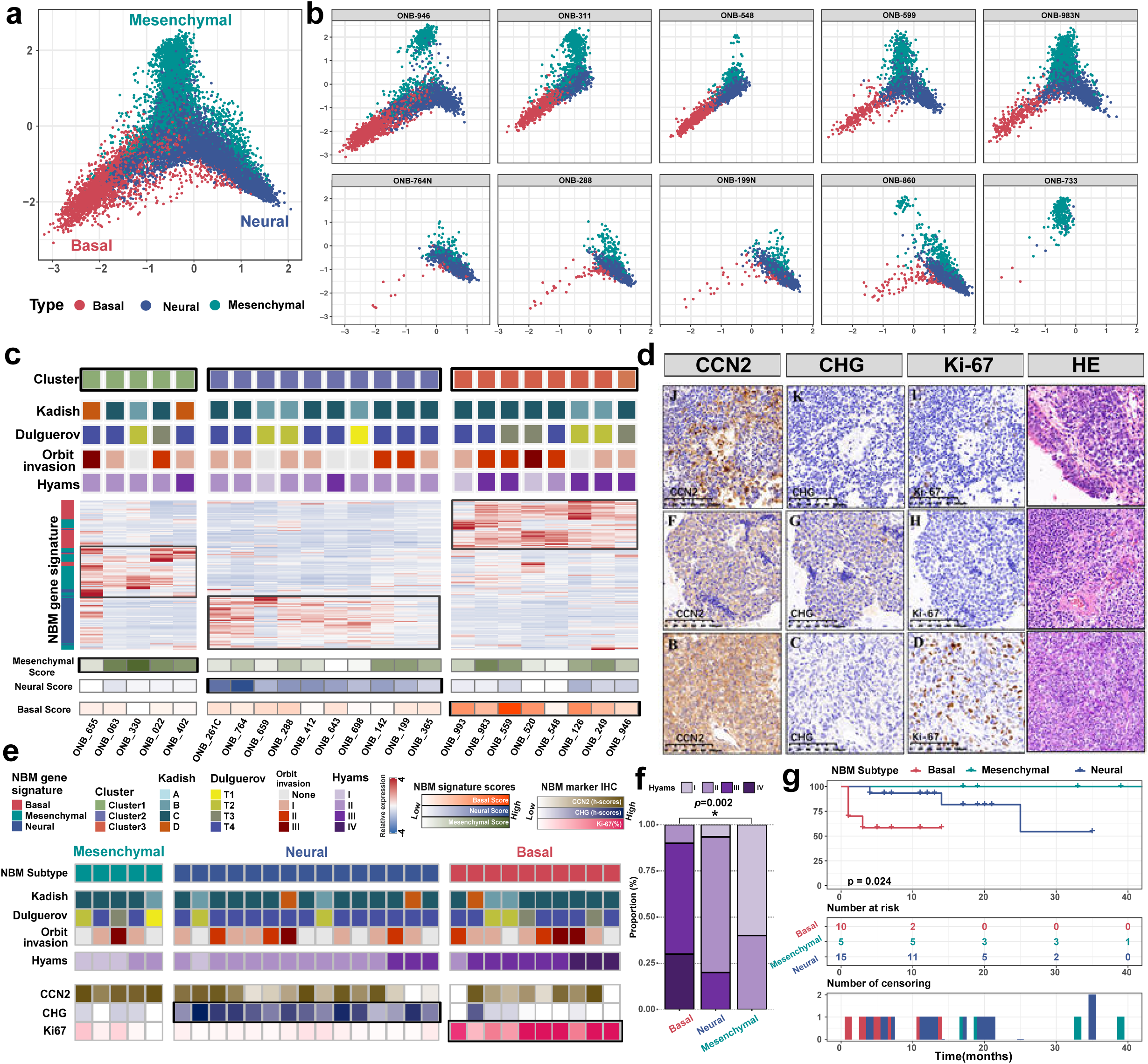
Characterization of three subtypes of ONBs based on scRNA-seq profiling. **a)** A ternary diagram depicting the NBM (Neural/Basal/Mesenchymal) Scores of each malignant single cell of ONB. **b)** Ternary diagrams showing the NBM Scores of malignant epithelial cells from each 10 ONB samples. **c)** Projection of scRNA-seq NBM malignant features onto bulk patient cohort stratified 23 ONB samples. The upper panel: The clinical characteristics of all 23 ONB patients, including modified Kadish stage, Dulguerov stage, Hyams grade, and grade of orbital invasion, were indicated as different colors. The middle panel: Heatmap showing expression score of each gene of NBM Top 50 signatures **(Table S8)** of each ONB patient in the bulk cohort. The bottom panel: the expression scores of Neural/Basal/Mesenchymal modules of each 23 ONB patients, scores evaluated (See **Material & Methods**). **d)** Representative immunohistochemistry (IHC) images of ONB samples showing HE staining results and the gradient of immunohistochemistry (IHC) scores of CHG, CCN2 and Ki-67(%) in 3 typical cases of ONB tumors, respectively. Scale bars, 100 µm (400x). **e)** IHC score-based subgroup prediction of 30 ONB patients with available paraffin section specimen. The upper panel: The clinical characteristics of all 30 ONB patients, including modified Kadish stage, Dulguerov stage, Hyams grade, and degree of orbital invasion, were indicated as different colors. The lower panel: relative immunohistochemistry-based expression levels of CHG (marker of Neural subtype), Ki-67% (marker of Basal subtype) and CCN2 (marker of Mesenchymal subtype). 30 ONB tumors were stratified into predicted NBM tumor subtypes based on expression patterns of representative markers. **f)** Bar graphs displaying comparisons of distributions of values for Hyams grade between tumor groups classified based on IHC score-based subgroup prediction. P-value was calculated using the Fisher exact test. *P < 0.05; **P < 0.01; ***P < 0.001; ns., no significance. **g)** Overall survival curves of a ONB cohort of 30 patients, stratified by NBM tumor groups classified based on Immunohistochemistry (IHC) score-based prediction (p = 0.024). P-value was calculated using log-rank test.

A bulk RNA-seq cohort consisting of 23 ONB patients receiving surgical resection at Eye, Ear, Nose and Throat Hospital of Fudan University was adopted as derivation cohort to further investigate the phenotypes of ONB. Using top 50 up-regulated DEGs as the classifier markers of Neural/Basal/Mesenchymal malignant cells **(Table S8)**, we conducted semi-supervised clustering, which divided 23 ONBs patients into 3 major groups **(Figure 3c)**. Group 2 and Group 3 were featured by their respective enrichment of Neural and Basal signatures, confirming the classification of classical Neural and Basal ONB subtypes. However, sorting the samples by their expression of NBM signatures revealed a group of tumors (Group1, N=5) harboring exclusively mesenchymal signatures. Furthermore, the bulk RNA-seq cohort containing 18 ONB patients provided by Classe et al. (GSE118995)^9^, based on which the researchers established the well-known binary classification of Basal/Neural ONB, were taken as an external validation cohort, and these patients also segregated into three clusters using NBM top 50 gene sets as signatures for semi-supervised clustering, similar to the three-class categories in our derivation cohort **(Figure S3a)**. Ultimately, the novel subtype of ONB was named as Mesenchymal due to its enrichment of mesenchymal gene signatures.

The Neural/Basal/Mesenchymal scoring systems established by scRNA analysis were projected onto bulk RNA-seq samples **(Figure 3c,** see **Materials & Methods)**. The expression patterns of NBM scores in the derivation cohort were found similar to the tripartite structure observed in scRNA ternary diagram. We noticed that most Basal ONB samples were also enriched for Neural and Mesenchymal Scores, and Neural subtype patients were could also be found with high Mesenchymal Scores, while in cases of Mesenchymal ONBs, tumors showed exclusively high Mesenchymal expression, which indicated that this novel ONB subtype is characterized by a double-negative expression of Neural/Basal features as well. We note that our findings also altered our knowledge of the classical Neural/Basal binary classifications, that determination of the Basal subtype takes priority over the Neural subtype.

In order to further validate our findings at protein expression level, Immunohistochemistry (IHC) was performed on a ONB cohort consisting of 30 patients to assess the proteomic expression of biomarkers representing NBM subtypes **(Figure 3d)**. Based on an overall consideration of relative abundance of DEGs across subtypes and frequently-used markers in previous studies, Ki-67, CHG (including CHGA and CHGB) and CCN2 (alias connective tissue growth factor, CTGF) were defined as markers of the Neural, Basal and Mesenchymal subtypes, respectively. The Basal group was identified with high proportion of Ki67 expression (≥25%) while the Neural group exhibited universally high level for CHG. We managed to confirm the existence of Mesenchymal ONBs by recognizing a few cases with absence of neither positive CHG expression nor high Ki-67% levels, but universal expression of CCN2 in tumor cells, consistent with our scRNA and bulk RNA-seq results **(Figure 3e)**.

Based on our findings validated by three expression platforms including single-cell RNA seq, bulk RNA-seq and IHC quantitative analysis, we established a flowchart to roughly stratify the 30 ONB samples into three subtypes using a relatively applicable approach in clinical practice **(Figure 3d, Figure S3b)**^22^. Tumors were first categorized by high Ki-67 level (Basal subtype if Ki-67%≥25%)^9^, hereafter by universally positive expression of Neural biomarker CHG (Neural subtype if H-score of CHG≥75), and then determined by their solely high expression of CCN2 (Mesenchymal subtype if Ki-67%<25%, H-score of CHG<75, H-score of CCN2≥75). Our findings suggested possible correlations between ONB subtypes and differentiation status of tumors. Mesenchymal, Neural and Basal ONBs exhibited a successively increased tendency of de-differentiation reflected by the Hyams grades (P = 0.002, Fisher’s exact test, **Figure 3f**). Survival analysis on disease free survival (DFS) stratified by NBM subtype also revealed statistical difference in prognostic outcomes among different subtypes of ONB with the Mesenchymal subtype showing the most optimistic survival results (P = 0.024, **Figure 3g)**.

### Depicting Cellular Diversity of Olfactory Lineages and Exploring the Cell-of-origin of ONB Subtypes

The olfactory mucosa (OM) has been known to maintain the capacity of constant neurogenesis in adulthood, which is sustained by continuous proliferation and differentiation of globose basal cells (GBCs). OM also retains the ability of regeneration after severe injury, by conditional activation of horizontal basal cells (HBCs) and almost all kinds of olfactory cell lineages could get reconstituted^12^. Based on the widely accepted hypothesis that ONB originates from normal olfactory epithelium, we firstly investigated the subclusters of normal olfactory lineages to further peruse the putative cell-of-origin of ONB subtypes, using a scRNA dataset of adult OM published by Durant et al (GSE139522)^11^.

Following protocols provided by original researchers, we gained N = 11348 cells comprising cellular components of adult olfactory epithelium **(Figure S4a)**. Cell types were assigned according to multiple known marker genes of human and murine OM **(Figure S4b, Table S9)**^11–13^. 6 clusters corresponding to olfactory stem cell lineages were identified, including HBCs (TP63, KRT5), GBCs (ASCL1, HES6), sustentacular cells (CXCL17, CYP2J2), mature and immature neuronal subgroups (iORNs and mORNs; GNG8, GNG13), as well as a population of immediate neuronal precursors (INPs, LHX2, NEUROD1). we performed a second round of clustering exclusively for these cells, which resulted in HBCs, GBCs, olfactory neuronal cells including INPs, iORNs, mORNs, and two sustentacular lineages consisting iSus and mSus defined by canonical markers for olfactory epithelium development **(Figure 4a-4b)**.

**Figure 4.**
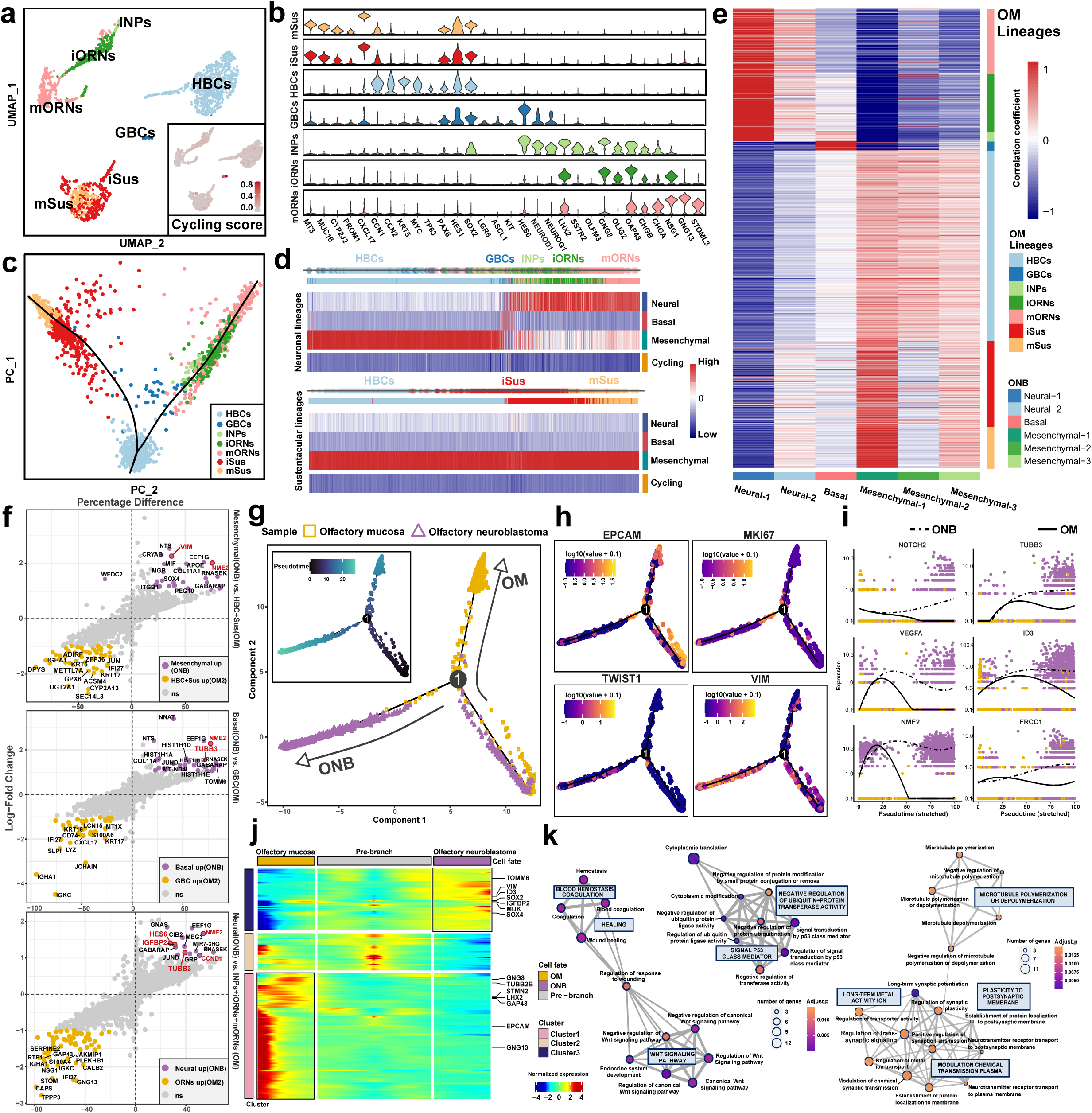
Exploring similarity and transcriptional diversity between olfactory epithelial lineages and ONB malignant cells. **a)** UMAP visualization showing the separation of the adult human olfactory lineages into HBCs, GBCs, INPs iORNs, mORNs and iSus, mSus. **b)** Panels of violin plot for expression levels of marker genes in olfactory neuronal and sustentacular lineages. **c)** A two-dimensional representation of single cell gene expression profiles based on principal component analysis. Slingshot predicted an early bifurcation in the lineage trajectories of the neuronal and sustentacular cells. **d)** Expression patterns of NBM malignant signatures in the olfactory neuronal and sustentacular lineages suggested similarities between ONB subtypes and diverse normal olfactory epithelial components. **e)** Heatmap showing Pearson correlations between scRNA expression profiles of ONB malignant subgroups (columns) and scRNA-seq profiles of olfactory epithelial cells (rows). Olfactory epithelial cells (rows) were arranged according to differentiation sequence of neuronal and sustentacular lineages. **f)** Scatter diagrams displaying the DEGs between Neural/Basal/Mesenchymal tumor cells and olfactory epithelial subclusters correspondingly related to these malignant subtypes (Neural vs INPs+iORNs+mORNs; Basal vs GBCs; Mesenchymal vs HBC+iSus+mSus). DEGs with Top 20 logFC were highlighted by vibrant colors (yellow & purple), specially expressed markers were labeled with gene symbol names. **g)** A pseudotime trajectory constructed using the combination of olfactory epithelial cells and malignant cells from a representative Basal type ONB (ONB-311). Different colors and shapes indicated the sample group of each cell (normal or tumor). **h)** Developmental trajectory of olfactory epithelial cells and malignant cells from ONB-311. Color gradient indicated the expression level of signature genes (EPCAM, MKI67, TWIST1, VIM). **i)** Pseudotime kinetics reflecting the varying expression levels of signature genes from the pre-branch of the trajectory to cell fate of normal OM (solid line) and ONB tumor (dashed line). **j)** Pseudo-heatmap displaying genes varying significantly along the pseudotime trajectory of olfactory epithelial cells and malignant cells from ONB-311. Pseudotime increased from the center to the left edge (olfactory mucosa, OM) and the right edge (olfactory neuroblastoma, ONB) of the heatmap. Rows are clustered and grouped by hierarchical clustering, with canonical transcription factors of olfactory neurogenesis and ONB tumorigenesis highlighted **(Table S11)**. **k)** Nerworks displaying clusters of representative upregulated GO pathways in olfactory neurogenesis and ONB tumorigenesis, respectively. Nodes indicate GO terms, with yellow color indicating smaller adjusted p value. The stronger the similarity between GO terms, the shorter and thicker the connecting edges. The size of the nodes indicated the number of genes contained in the GO pathway. HBC, horizontal basal cell; GBC, globose basal cell; INP, immature neuron progenitors; iOSN, immature olfactory sensory neuron; mOSN, mature olfactory sensory neuron; mSUS, mature sustentacular cell; iSUS, immature sustentacular cell. DEGs, differentially expressed genes.

Slingshot trajectory analysis, a statistical framework for cell fate tracing proposed by Fletcher et al. to elucidate complex cell differentiation in murine olfactory neurogenesis, was applied in our study to display olfactory lineage trajectories in adult human OM **(Figure 4c)**^12^. We established a dichotomous structure in which the sustentacular cells developed through direct fate conversion from HBCs, whereas the neuronal lineages expanded via proliferation of GBC and INP progenitors, resembling the developmental paths observed in murine OM.

Next, in order to search for the diverse origins of ONB subtypes, we compared the transcriptomic profiles between neural/basal/mesenchymal subpopulations and cells in adult olfactory epithelium. We created single dimensional plots to explore the average expression of cycling and NBM signatures varying according to the developmental distance of cells within olfactory neuronal and sustentacular lineages, respectively **(Figure 4d)**. GBCs were proved to be the proliferative progenitors in normal adult OM. We discovered that GBCs committed Basal subtype features, and enrichment of Neural subtype signatures were found in olfactory neuronal lineages (INPs, iORNs and mORNs). Unexpectedly, gene signatures of the novel mesenchymal subtype were observed to be enriched in the sustentacular lineages, which corroborated with results of evaluating expression scores of NBM signatures in olfactory lineages using the Seurat function AddModuleScore() **(Figure S4c)**. Then we performed Pearson’s correlation a of adult human olfactory cells and ONB malignant subgroups except the cilium cluster, which showed that the Basal malignant cells resembled GBCs, and revealed links between the Neural subgroups and neuronal olfactory linages ranging from INPs, iORNs to mORNs. We also discovered similarities between the Mesenchymal subpopulations and sustentacular lineages, including HBCs and two sustentacular clusters especially, which were regarded as direct differentiation products of HBCs that does not require cell division.

### Molecular Features Underlying Malignant Transition from Normal OM to ONB

To demonstrate the transcriptional differences between normal olfactory lineages and ONB malignant cells, we performed differential expression analysis comparing NBM malignant subpopulations and their corresponding suspected normal cell origins, including Basal tumoral cells and GBCs, Neural malignant cells and neuronal olfactory lineages (INPs, iORNs and mORNs), Mesenchymal tumor cells and sustentacular lineages (HBCs, iSus, and Sus) **(Figure 4f)**. In comparison to normal olfactory epithelial, ONB cells were predominantly characterized by upregulated expression of markers related to neurogenesis and tumor promotion, including TUBB3 (encoding protein TUJ1), HES6, VIM and MNE2, etc **(Table S10)**. The findings suggested that despite an evident existence of developmental relationship between normal olfactory lineages and ONB tumor cells, the cancer cells have acquired unique expression patterns underpinning malignant transformation, which could possibly shed light on discovery of therapeutic targets and diagnostic markers for this unusual pathology. For instance, strong expression of TUBB3, which is also a marker for vincristine resistance, were confirmed in 28/30 ONB patients through IHC staining. Moreover, we found low expression of TUBB3 in 10/12 sinonasal melanoma including one patient who had been misclassified as ONB by preoperative pathological diagnosis **(Figure S4d-S4e)**. TUBB3 might be proposed as a diagnostic marker for ONB that could possibly distinguish this unique malignancy from other small round blue cell tumors of the skull base, which contains several assorted neoplasms that often get pathologically misdiagnosed from one another.

We integrated normal olfactory lineages (OM2) with malignant cells of ONB patient (ONB-311) and conducted pseudo-time analysis, which resulted in a bifurcated differentiation trajectory emanating from HBCs, with one branch representing normal development towards olfactory neurogenesis and the other comprised predominantly of ONB tumor cells **(Figure 4g, Figure S4f)**. Cells with proliferative expression were positioned at the pivot of bifurcation in the developmental trajectory **(Figure S4g)**. Our results confirmed the developmental relationship between ONB and normal olfactory epithelium at single cell resolution.

We then explored the plasticity and transcriptional features underneath malignant transition of ONB. Tumor cells exhibited high expression levels for molecules related to EMT (VIM and TWIST), angiogenesis (VEGFA) and NOTCH pathway, as well as decreased expression of EPCAM. Increased expression of ERCC1 (marker for cisplatin resistance in non-small cell lung cancer) and NME2 (marker for 5-FU resistance) revealed latent targets that might facilitate treatment decision **(Figure 4h-4i)**. Dynamic expression changes along the trajectory of olfactory development and ONB malignant transition were displayed by pseudo-heatmap **(Figure 4j and Table S11).** Functional enrichments of gene clusters preferentially expressed in tumoral and normal cell fates were analyzed using GO/KEGG pathway analysis **(Figure 4k)**. Compared with normal olfactory epithelium, tumor cells were enriched for signaling pathways such as blood coagulation wound healing, P53 class mediator, ubiquitin-protein transferase activity, and WNT signaling pathway.

### Diversity and dynamics of T cell subpopulations in ONB tumor microenvironment (TME)

Tumor-infiltrating T cells (TILs) have been widely known to play a central role in the TME, dominating critical clinical properties so as immunotherapy-related responses^23^. Re-clustering of N = 26996 NK/T cells revealed 3 major classifications including 5 subclusters of CD4^+^ T, 5 subclusters of CD8^+^ T, 2 subclusters of NK cells and an undefined subgroup consisting of low-quality cells **(Figure 5a)**. We found that CD8-GZMH, CD8-PDCD1 and NK-FGFBP2 cells distributed predominantly in tumor TME **(Figure 5a)**. Referring to reported T cell function-associated gene sets **(Figure 5b&5c)**^24, 25^, we managed to annotate the identities of several clusters of CD8^+^ T cells (GZMH-effector, GZMK-effector memory, ZNF683-resident memory and PDCD1-exhausted), and conventional CD4^+^ T cells (CCR7-naïve, IL7R-central memory and IL21-follicular helper), as well as FOXP3-resting and CTLA4-immuno-suppressive populations of CD4^+^ Tregs^26^.

**Figure 5.**
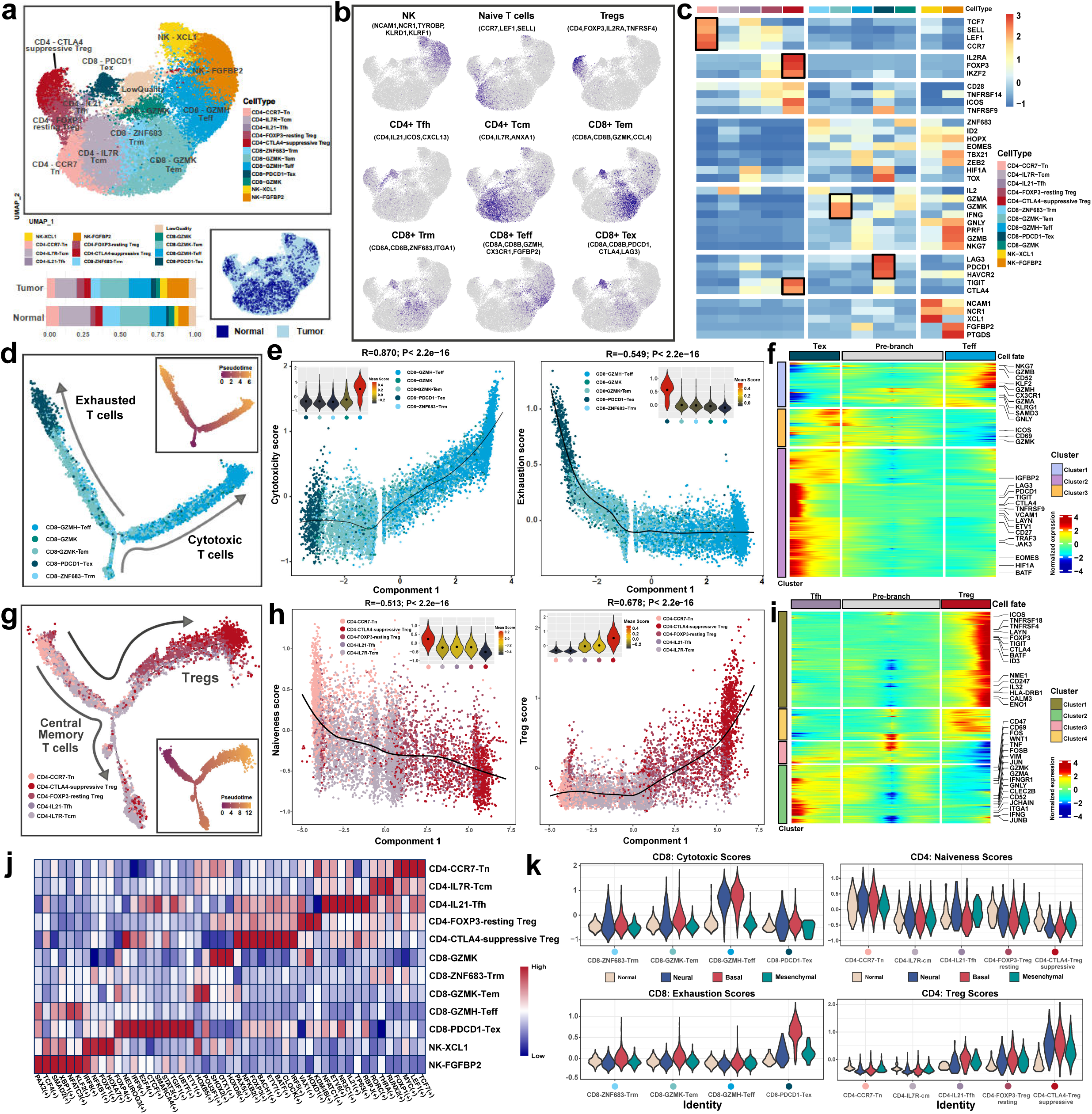
Identification of diverse functional status and subtypes of T cells in ONB tumor microenvironment. **a)** UMAP visualization of 26996 T/NK cells showing the formation of five CD4^+^ and five CD8^+^ T cell subtypes, and two NK cell subtypes, colored by cell type (top) and different tissue origins (bottom right) and stratified by proportions of cell types in normal and tumor tissues (bottom left). **b)** UMAP visualization, colors representing the expression levels of canonical markers (gray to purple) for the indicated T cell subtypes. **c)** Z-score normalized mean expression of selected T cell function-associated genes in each cell cluster. Black boxes highlight the prominent patterns defining known T cell subtypes. **d)** Inferred differentiation trajectory of CD8^+^ T cells. Arrows showed the developmental directions of CD8^+^ T cell properties (Tex & Teff) annotated with the signatures shown in **b & c**. **e)** Traceplots depicting varying expression levels of functional features of CD8^+^ T cells (Exhaustion Scores & Cytotoxicity Scores) along Monocle2 components, calculated by the mean expression of gene sets related to these T cell functions. The solid lines represent LOESS fitting of the relationship between these scores with Monocle2 components. Violin plots (top corners) showed the distribution of functional scores in CD8^+^ T cell clusters, colored by means of the corresponding scores in each cluster, with black dots representing median values. P values were calculated by Pearson correlation, and P < 2.2 × 10^-16^ represents a *p* value approaching 0. **f)** Pseudo-heatmap showing relative expression levels of canonical markers of CD8^+^ T cells along inferred trajectories. The dark-green and indigo-blue branches reflected the two developmental directions (Tex and Teff) in **e**, respectively. **g)** Inferred differentiation trajectory of CD4^+^ T cells. Arrows showed the developmental directions of CD4^+^ T cell properties (Tcm & Tregs) annotated with the signatures shown in **b & c**. **h)** Traceplots depicting varying expression levels of functional features of CD4^+^ T cells (Naiveness Scores & Treg Scores) along Monocle2 components, calculated by the mean expression of gene sets related to these T cell functions. Violin plots (top corners) showed the distribution of functional scores in CD4^+^ T cell clusters. P values were calculated by Pearson correlation. **i)** Pseudo-heatmap showing relative expression levels of canonical markers of CD4^+^ T cells along inferred trajectories. The purple and red branches reflected the two developmental directions (Tcm & Tregs) in **g**, respectively. **j)** Heatmap showing the activity of TFs in each T/NK cell subtype. The TF activity is scored using AUCell. **k)** Violin plots showing the distribution of Exhaustion Scores & Cytotoxicity Scores in 5 CD8^+^ T cell subclusters, and Naiveness Scores & Treg Scores in 5 CD8^+^ T cell subclusters, stratified by NBM (Neural/Basal/Mesenchymal) tumor subtypes of ONB.

Pseudotime trajectory analysis by Monocle2 revealed a dichotomous structure of both CD4^+^ T and CD8^+^ T cell differentiation, which reflected the classical perspective on T cell developmental dynamics proposed by previous studies **(Figure 5d&5g)**^24, 27, 28^. Owing to the absence of naïve-like CD8^+^ T cells found in the ONB TME **(Figure 5c)**, the dichotomy started from CD8-GZMK-Tem and CD8-ZNF683-Trm, and then developed into either cytotoxic (CD8-GZMK-Teff) or dysfunctional (CD8-PDCD1-Tex) T cells. Consistent with previous findings, we identified the effector memory (CD8-GZMK) T cells in ONB, which presented with a pre-dysfunctional status, characterized by both high expression level of GZMK and intermediate expression of PDCD1, LAG3 (**Figure S5a)**^29^. The exhausted CD8-PDCD1-Tex cells at the terminal of inferred trajectory were found with the highest pseudotime scores, though presented diluted cytotoxicity as activated T cells, which indicated dysfunction induced by tumoral immune response in ONB **(Figure 5e, Figure S5b, Table S12)**. The transition of expression patterns in CD8^+^ T cells in ONB were further displayed in the pseudo-heatmap **(Figure 5f, Table S13)**. In addition to immunosuppressive receptors including LAG3, CTLA4 and PDCD1, the exhausted CD8^+^ T cells also expressed pro-angiogenesis (HIF1A) and immuno-regulation (TIGIT) features^30–32^.

The activated-exhausted coupled intrinsic of TILs in ONB was further revealed by evaluating Spearman’s correlation between CD8 T cell activity, which was represented by the average granzyme expression (GZMA, GZMB and GZMH), and expression of 233 CD8 T-cell specific genes **(Figure S5c, Table S14)**^33^. LAG3 and TIGIT were found positively associated with T cell activity, and relative enrichment of CD8^+^LAG3^+^ T cells in tumor stroma compared to peri-tumoral mucosa has been confirmed by immunofluorescence (IF) staining **(Figure S5d)**, which suggested LAG3 as potentially effective immunotherapy target for ONB patients.

The branched developmental trajectory of CD4+ T cells initiated from naïve CD4^+^ T (CD4-CCR7-Tn) and bifurcated toward either follicular helper T cells (CD4-IL21-Tfh) or to an immune regulatory phenotype (Treg-FOXP3-resting and Treg-CTLA4-suppressive) **(Figure 5g)**. The CTLA4-suppressive Tregs represented a mature subcluster with the lowest naiveness and the highest Tregs scores **(Figure h, Table S15)**. The continuously increasing expression spectrum of immuno-suppressive and co-stimulatory features including CTLA4, LAYN, BATF and TNFRSFs reflected a gradual activation of Tregs in ONB **(Figure 5i, Figure S5e)**.

SCENIC analysis was performed to investigate the transcription networks between diverse T cell subtypes **(Figure 5j)**. Transcription factors (TFs) such as IRF9, which has been reported to induce PDCD1^34^, and ETV1 that targeting exhaustion marker TOX^35^, were found with high regulon specificity in the Tex subgroup, suggesting the latent role of these TFs to trigger transition toward exhausted state of CD8^+^ T cells. We also observed an elevated transcriptional activity of NFATC1 that was reported to regulate PD-1 expression^36^, in CD4-IL21-Tfh cells, suggesting that Tfh cells in ONB might assist T cell dysfunction **(Figure S5f)**.

Then we compared the Cytotoxic & Exhaustion Scores of CD8^+^ and Naiveness & Treg Scores of CD4^+^ T cells across tumor-infiltrating and normal tissue-derived T cells **(Figure 5k, Figure S5g)**. TILs from Basal and Neural subtype of ONBs tumors harbored increased Cytotoxicity Score, while TILs from Basal subtype showed high Exhaustion Scores exclusively, indicating that patients with Basal subtype ONB might possibly acquire higher response rate to immune checkpoint blockers (ICBs).

### Depicting Tumor-associated Myeloid cell Phenotypes

To delineate the diversity of tumor infiltrating myeloid lineages in ONB, sub-clustering of N = 14951 myeloid cells were performed. 5 categories consisting of 14 subpopulations **(Figure 6a)**, including mast cells (Mast-CPA3), dendritic cells (DC-CD1C, DC-CLEC9A, DC-LAMP3), macrophages (Macro-C1QA, Macro-CD163, Macro-GPNMB, Macro-RGS1, Macro-SPP1), and neutrophils (Neu-CSF3R, IFN-response), were annotated based on classical myeloid markers **(Figure 6b)**. The cellular proportions of diverse myeloid subgroups exhibited disparities between normal olfactory and ONB tumoral tissue origins **(Figure 6a, Figure S6a)**. GSVA pathway enrichment analysis uncovered diverse functional roles played by tumor-associated macrophages (TAMs), including complement activation in Macro-C1QA, antigen presenting and immune cell recruitment which stands for pro-inflammatory response in Macro-RGS1. Notably, up-regulated hypoxia, angiogenesis, extracellular matrix formation related pathways were observed in Macro-SPP1, suggesting tissue-remodeling and pro-angiogenic activity associated with tumoral promotion **(Figure 6g)**.

**Figure 6.**
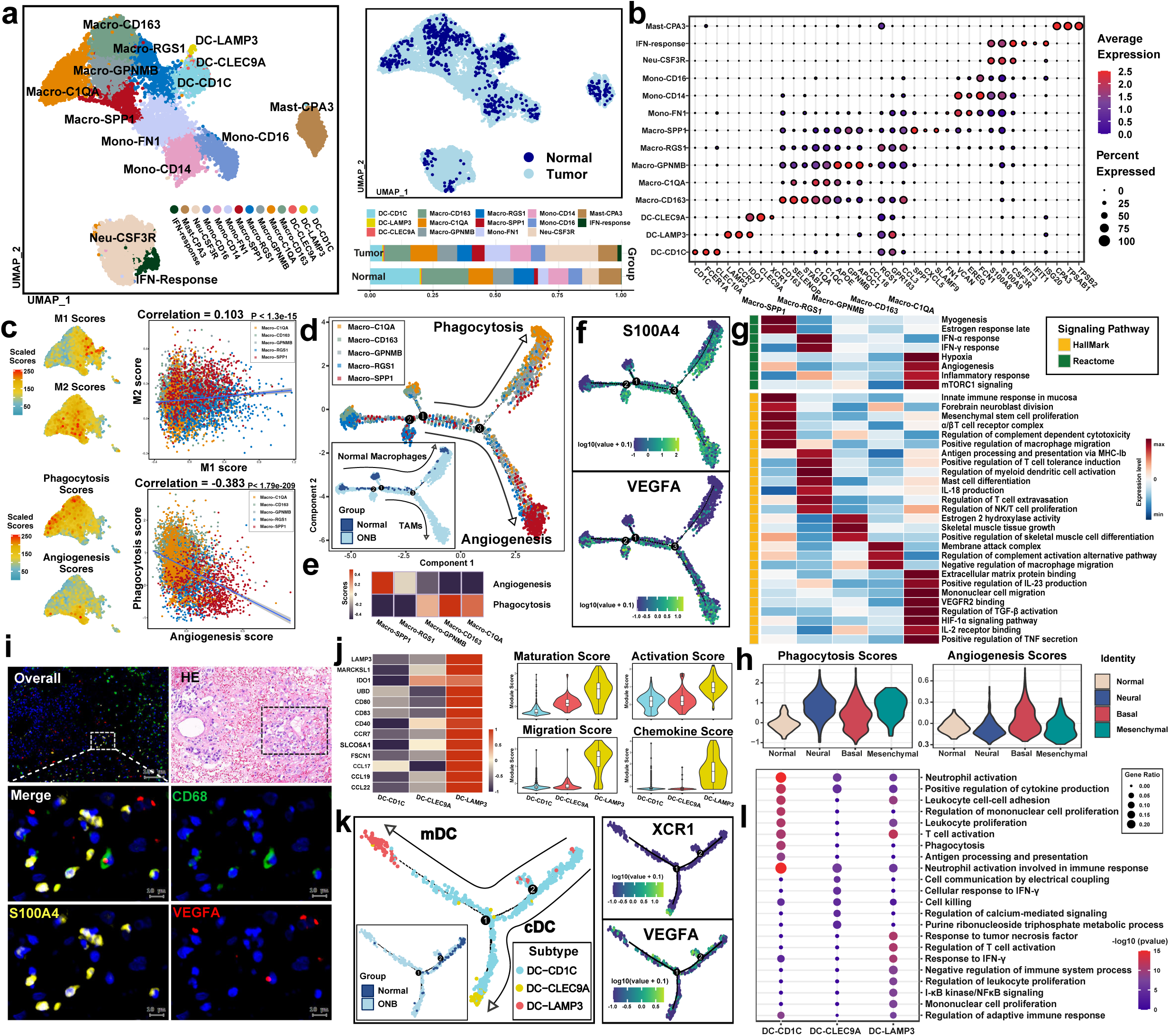
Characterization of multiple functions and subclusters of myeloid cells in ONB tumor microenvironment. **a)** UMAP visualization of 14951 myeloid cells showing the formation of 3 DC subclusters, 5 macrophage subclusters, 3 monocyte subclusters and a cluster of mast cells, colored by cell type (left) and tissue type (right top) and stratified by proportions of cell types in normal and tumor tissues (right bottom). **b)** Violin plots showing the expression levels of different marker genes across 14 myeloid subclusters. **c)** UMAP plots displaying expression levels of known signatures indicating functional status of myeloid cells (M1 & M2 Scores, Phagocytosis & Angiogenesis Scores) in macrophages. Scatterplot showing the Pearson correlation between M1 & M2 Scores, Phagocytosis & Angiogenesis Scores. **d)** Inferred developmental trajectory of macrophages. Arrows in the main plot revealed a dichotomic functional division (Phagocytosis & Angiogenesis). The two branches were predominantly comprised of macrophages originating from normal olfactory epithelium or ONB tumor microenvironment, respectively. **e)** Heatmap showing different expression patterns of phagocytosis- and angiogenesis-associated signature genes among macrophage subsets in ONB. **f)** Developmental trajectory of macrophages. Color gradient indicated the expression level of signature genes. **g)** Heatmap showing the selected signaling pathways that were significantly enriched in GSVA analyses for each macrophage cell subcluster derived from GO and Reactome signaling pathways. Filled colors from dark-blue to red represent scaled expression levels (normalized −log10 p values) from low to high. **h)** Violin plots showing the distribution of Phagocytosis & Angiogenesis Scores of macrophages, stratified by tissue origins including normal OM and NBM (Neural/Basal/Mesenchymal) tumor subtypes of ONB. **i)** Representative images of multiplex immunofluorescence (IF) staining of macrophages in ONB tissues. Proteins detected using respective antibodies in the assays are indicated on top. Scale bars, 10 µm. **j)** Heatmap and violin plots exhibiting the module scores and gene expression levels of functional-related gene sets (maturation, activation, migration, and chemokine ligand) in 3 DC subclusters. Box plots inside the violins indicated the quartiles of corresponding score levels. **k)** Inferred developmental trajectory of DCs (left). Arrows in the main plot reflected a dichotomic division (Phagocytosis & Angiogenesis). The root and branches of the trajectory were predominantly comprised of DCs originating from normal olfactory epithelium or ONB tumor microenvironment, respectively. Trajectory displayed with color gradient (right top & right bottom) indicated the expression level of canonical signature genes for DC subtypes. **l)** Dot plot showing the selected signaling pathways that significantly enriched in GO signaling pathways for each DC subcluster.

The classical and alternative activation of macrophages, namely M1/M2 polarization, have long been featured by distinct transcriptome patterns, such as CD86 and tumor necrosis factors (TNFs) for M1, CD163 and matrix metalloproteinases (MMPs) for M2 **(Table S16)**^37, 38^. Generally, M1-like macrophages with pro-inflammatory cytokine secretion present as facilitators of tumor-suppressive response. In contrast, M2-polarization of TAMs were often regarded as transition toward a tolerogenic, anti-inflammatory phenotype^39, 40^. Zhang et al.^41^ described functional division of TAMs into phagocytic and pro-angiogenic phenotypes, exhibiting a dichotomous activation model distinctive from M1/M2 classifications.

High level expression of M1 and M2 features were respectively observed in Macro-RGS1 and Macro-SPP1 subclusters, while most macrophages showed increased expression while no obvious preference of M1/M2 Scores, suggesting a coupled M1-M2 activation status **(Figure 6c)**^23^, as previously observed in TAMs of nasopharyngeal carcinoma^24^. Pearson correlation analysis revealed significantly negative relations between Phagocytosis/Angiogenesis Scores **(Figure 6c)**. Moreover, trajectory analysis revealed a dichotomous differentiation of phagocytic and pro-angiogenic phenotypes, with macrophages from normal olfactory mucosa located mainly along the differentiation branch of phagocytosis **(Figure 6d)**. Macro-C1QA was supposed to function as the phagocytic cluster with a terminally differentiated status and increased expression level of phagocytic features. In the contrary, Macro-SPP1 was identified as a typical TAM subgroup in ONB, with predominantly tumoral tissue origin and exclusive enrichment for regulators of angiogenesis and immunosuppressive, marked by VEGFA and SPP1. **(Figure 6e-6f,** Figure S6b-S6c) We found elevated expression of S100A4 in macrophages derived from ONB **(Figure 6f, Figure S6d)**, a marker also known as metastasin which has been recently identified to enhance protumor (M2) polarization in TAMs^42^. The existence of S100A4^+^ and VEGFA^+^ TAMs in microenvironment of ONB were confirmed by immunofluorescence assay **(Figure 6i, Figure S6e)**, suggesting an overlap of M2-tolerogenic and pro-angiogenic functional status. Inducing repolarization from M2-like protumor macrophages towards M1-like pro-inflammatory phenotype has become an active area for investigation to reverse immuno-suppression^43^. TAMs promote tumor growth by regulating blood vessel sprouting and remodeling, providing potential therapy targets for anti-tumor inhibitors^44^.

Based on the Neural/Basal/Mesenchymal (NBM) classification described previously in this study, macrophages were divided into 4 groups that from different sample subtypes (Normal OM and N/B/M subtypes of ONB). Macrophages from tumor samples were observed with both higher M1/M2 and Phagocytosis/Angiogenesis Scores when compared to that from normal olfactory mucosa, indicating multiple functional activation of TAMs **(Figure S6f)**. Of note, macrophages from Basal ONBs exhibited a tendency toward lower level of phagocytic activation and elevated expression of pro-angiogenic markers **(Figure 6h)**. The heterogeneity among ONB subtypes might help identify candidates who were more likely to benefit from targeted anti-angiogenesis therapy.

As a major population of professional antigen-presenting cells (APCs), dendritic cells have aroused interest for cancer immunotherapy due to their role in inducing protective adaptive immunity with antitumor effect^45^. We identified 3 subgroups of DCs, including cDC1 (CD1C^+^, XCR1^+^ and BATF3^+^) playing as efficient antigen cross-presenters to CD8^+^ T cells and cDC2 (FCRR1A^+^) which preferentially interacted with CD4^+^ T cells^46^. Notably, DC-LAMP3 were identified as the mDC (LAMP3^+^, CCR7^+^) cluster, a mature subtype expressing elevated levels of gene modules related to chemokines for immune cell recruitment, and DC maturation, activation, migration, consistent with previous studies **(Figure 6j)**^23, 47^. Migratory DCs were reported to bear the capacity to transfer antigen to lymph node-resident T cells to initiate immune response^48^. Pseudotime analysis revealed that DC-CD1C (cDC1) cells developed into DC-FCER1A (cDC2) and DC-LAMP3 (mDC) cells, and mDCs appeared to be the most differentiated DCs with highest inferred pseudotime score. DCs in ONB exhibited a continuous spectrum of activation states in the trajectory, while DCs from normal OM occupied mainly the pre-branch before dichotomy **(Figure 6k)**. GO and GSVA signaling pathway enrichment analysis uncovered diverse functional roles of DCs, addressing the role of antigen presenters of this active multi-tasking myeloid subpopulation **(Figure 6l, Figure 6Sg)**.

### Heterogeneity and functional diversity of stromal cell lineages in ONB

A total of N = 11698 stromal cells were extracted for re-clustering, in order to depict cellular heterogeneity of stromal lineages in ONB microenvironment **(Figure 7a)**. N = 5886 endothelial cells (ECs, marked by PECAM1 and PLVAP) was divided into 5 subgroups, and N = 3246 fibroblasts (COL1A1, COL1A2) consisted of 4 subclusters. Besides, a cluster expressing cell-cycling signatures (MKI67, TOP2A), as well as a cluster recognized as pericytes (RGS5), were respectively identified according to canonical markers **(Figure 7b)**.

**Figure 7.**
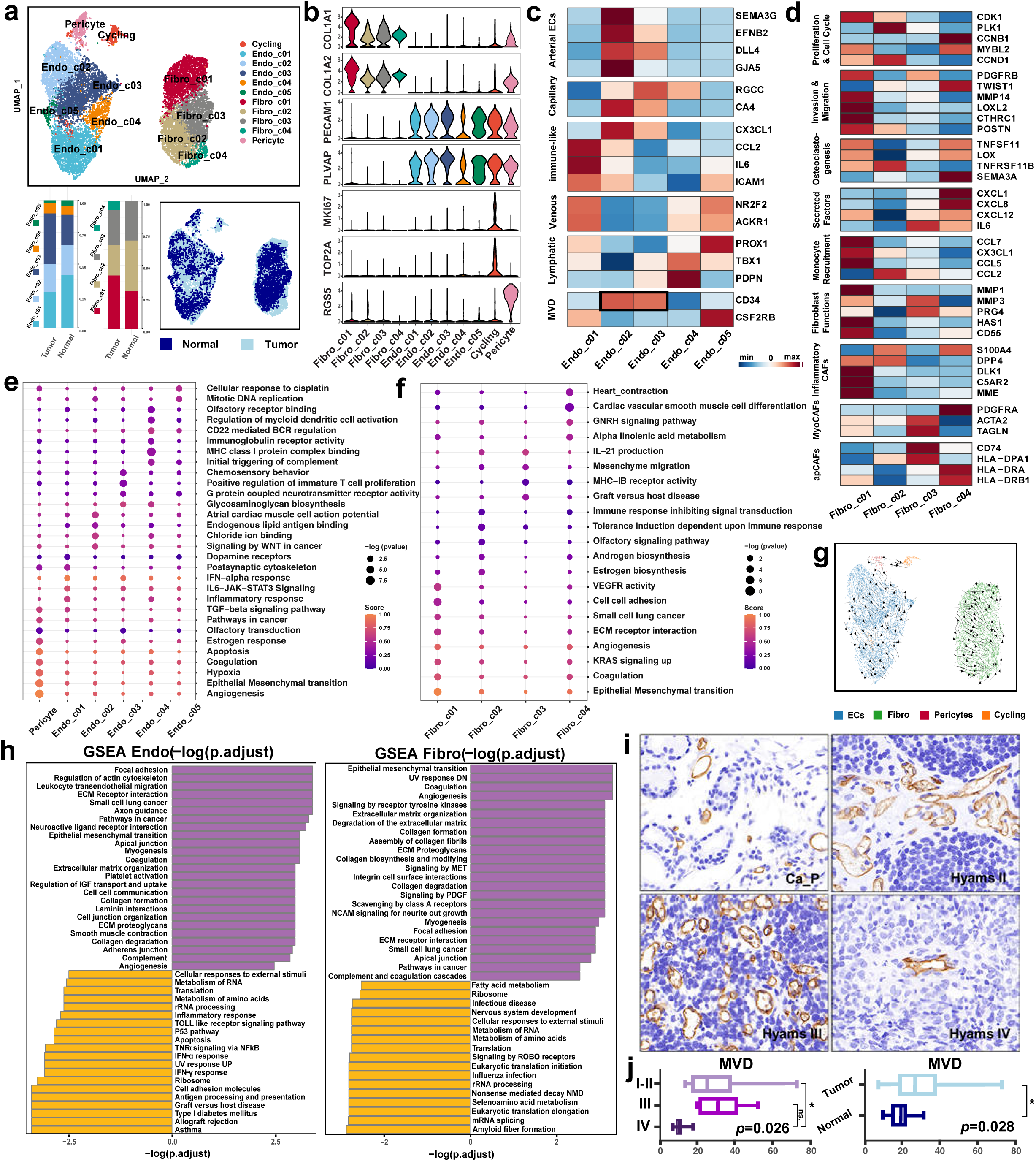
Cellular heterogeneity of ECs and dynamic functional states of fibroblasts in ONB tumor microenvironment. **a)** UMAP visualization of 11698 stroma cells showing the formation of 5 EC subclusters, 4 fibroblast subclusters, a cluster of pericytes, and a subset of cycling stromal cells, colored by cell type (top) and tissue type (bottom right) and stratified by proportions of cell types in normal and tumor tissues (bottom left). **b)** Violin plots displaying expression levels of marker genes in stromal subclusters in ONB tumor microenvironment. **c)** Heatmap showing expression levels of canonical marker genes of known EC subtypes in each EC subcluster. **d)** Heatmap exhibiting the expression levels of gene sets corresponding to canonical fibroblast subtypes and stromal cell functions in 4 fibroblast subclusters. **e)-f)** Dot plots displaying representative signaling pathways that were significantly enriched in GSVA analyses for pericyte and 5 EC subclusters **(e)** and 4 fibroblast subclusters **(f)** derived from GO, KEGG and Reactome signaling pathways. g) The RNA velocity trajectory and the latent time stromal cells in ONB microenvironment. h) Differences in pathway activities scored by GSEA analysis between malignant and non-malignant tissue derived ECs (left) and fibroblasts (right). i) Selected IHC staining images of CD34 which marked MVD in peri-cancerous mucosa and representative ONB tumor samples with different hyams grade (II, III and IV). Scale bars, 100 μm, 40×. g) Boxplots comparing MVD levels among Hyams II, III and IV grade ONB tumors, and between malignant and non-malignant tissue, respectively. The one-way ANOVA test was applied to compare MVD levels between malignant and non-malignant tissue. *P < 0.05; **P < 0.01; ***P < 0.001; ns., no significance.

**Figure 8.**
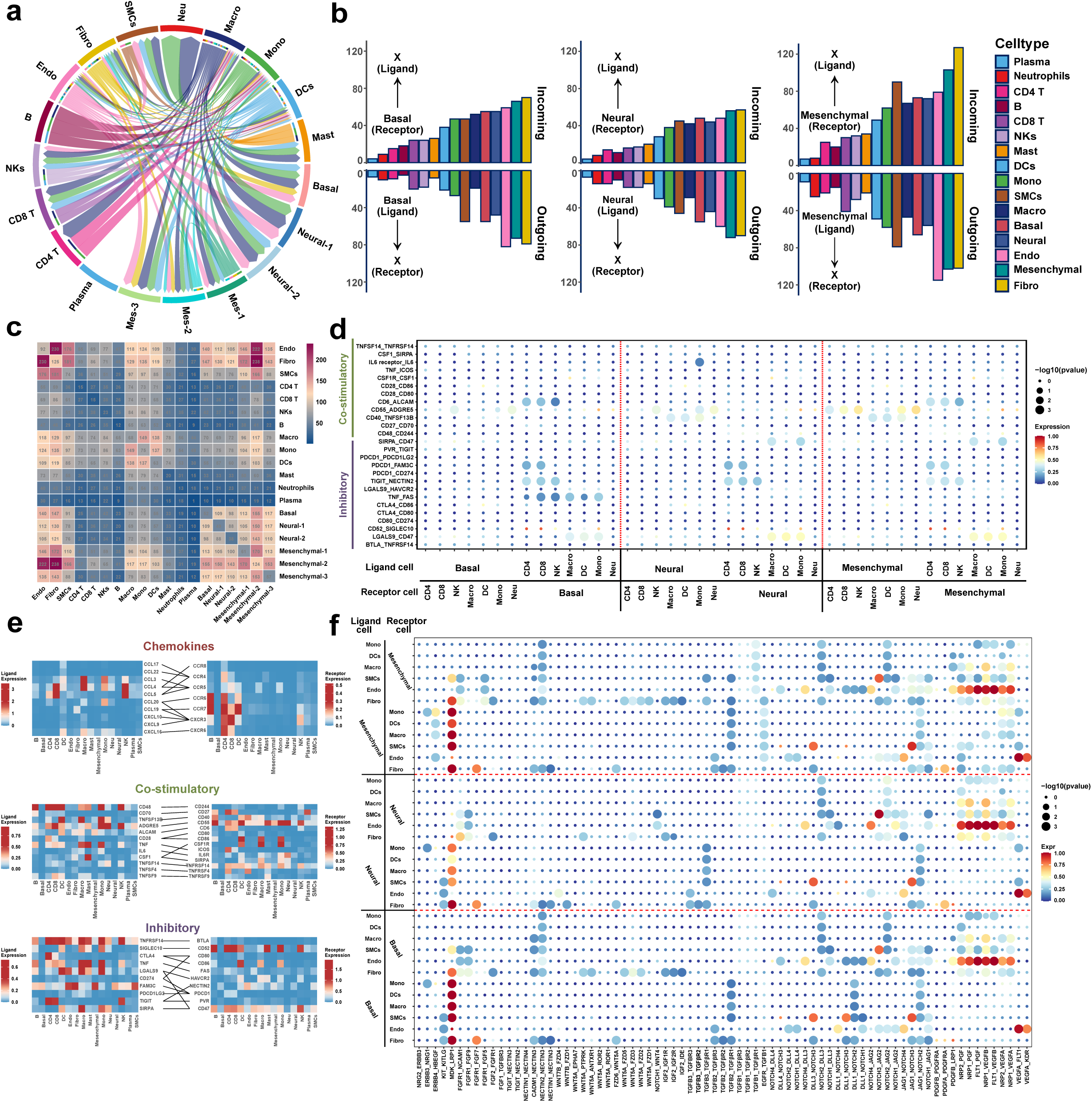
Intercellular interactions in ONB tumor microenvironment. a) Circo-plot showing the intercellular interactions among different cell types in ONB tumor microenvironment. The strings represented interactions determined on the basis of expression of a ligand by one cell type and expression of a corresponding receptor by another cell type. The thickness of strings reflected the amount of different interaction pairs, colored according to cell type. b) Bar plots depicting numbers of putative receptor-ligand interactions between Neural, Basal and Mesenchymal ONB malignant cells and indicated cell types. Interaction numbers were calculated based on expression of receptors and corresponding ligands in scRNA-seq data. Outgoing interactions refer to the sum of ligands from malignant cells that interact with receptors on the indicated cell type. Incoming interactions refer to the opposite. c) Heatmap showing numbers of potential ligand-receptor pairs among different cell types in ONB tumor microenvironment predicted by CellphoneDB. d) Dot plots showing selected ligand-receptor interactions between immune cells and malignant cell in ONB, separated by Basal (right), Neural (middle) and Mesenchymal (left) tumor subtypes. e) Heatmaps showing ligand-receptor interactions among different cell types in ONB tumor microenvironment. f) Dot plots showing mesenchymal ligand-receptor interactions between stromal cells, myeloid cells and malignant cell in ONB, separated by Mesenchymal (top), Neural (middle) and Basal (bottom) tumor subtypes.

ECs have been known to associate with tumor growth and progression, thus have increasingly become potential targets of anti-cancer therapies^49^. Identities of EC subclusters were annotated by assessing the expression patterns of distinctive markers in published studies **(Figure 7c)**^50^. Endo_c02-GJA4 and Endo_c03-COL4A1 shared a transcriptomic feature of capillary ECs (RGCC and CA4), while Endo_c02-GJA4 also expressed markers of Arterial ECs (SEMA3G, EFNB2, DLL4 and GJA5). Endo_c01-ACKR1 was classified as venous ECs for expression of venous marker NR2F2 and ACKR1. Endo_c04-MARCKSL1 was identified as lymphatic ECs (TBX1, PDPN), whereas Endo_c05-CD74, a small subset of capillary-like ECs, was enriched for antigen-presentation and immune-related transcripts (CX3CL1, CCL2 and ICAM1), was regarded as immune ECs. ECs in ONB were discovered to involve in many biological processes using GSVA analysis, including inflammatory response, immune-regulation, biological metabolism and even olfactory transduction. Notably, GSVA pathway analysis revealed that pericytes contributed to pathways in cancer, EMT, hypoxia and angiogenesis, suggesting the role of promoting tumorigenesis for this subpopulation **(Figure 7e)**.

Fibroblasts, featured by abundant cellular burden, highly heterogeneous functions and mutable expression patterns, have long been regarded as major players in tumor microenvironment to promote oncogenesis^51^. Fibro_c01-COL1A1 cells comprised larger cellular proportion in tumor-associated fibroblasts (CAFs), and Fibro_c04-C7 cells distributed exclusively in tumor stroma (**Figure 7a, Figure S7a**). We investigated expression patterns of functional molecules in each fibroblast subcluster **(Figure 7d)**^52^. The results showed that Fibro_c01 cluster marked diverse characteristics of activated CAFs and demonstrated multipotency corresponding to Monocyte recruitment, invasion and migration. Fibro_c01 cells exhibited transcriptomic features of both inflammatory CAFs (iCAFs, DLK1, C5AR2 and MME, **Figure 7d**) and myofibroblastic CAFs (myoCAFs, CTHRC1 and MMP11, **Figure S7b**)^53^. Fibro_c04-C7 cells showed up-regulated expression of molecules related to chemokine secretion (CXCL1 and CXCL8) and antigen presenting (HLA-DRA and HLA-DRB1). The terminal developmental state inferred by RNA velocity suggested that Fibro_c04-C7 was probably converted from CAFs induced by tumorigenesis to participate in tumoral Immune response **(Figure 7g)**. GSVA pathway analysis further revealed multiple roles including pro-angiogenesis and epithelial mesenchymal transition played by Fibro_c01-COL1A1 cluster. Hormone metabolic pathways, including androgen and estrogen biosynthesis, were found enriched in Fibro_c02-CPE. Fibro_c03-APOD was possibly regarded as antigen-presenting CAFs (apCAFs), defined by the co-expression of the invariant chain of MHC II molecule CD74 with HLA-DPA1^53^, which was further supported by GSVA analysis that indicated MHC-IB receptor activity of this fibroblast subset **(Figure 7f)**.

GSEA enrichment analysis was conducted to reveal functional diversity of ECs and fibroblast subclusters in normal olfactory mucosa and tumor TME^33^. The results both showed intensified extracellular matrix remodeling process and immune regulation, as well as angiogenesis in stromal cells from ONB **(Figure 7h)**. Since tumoral vessel sprouting has become an emerging therapeutic target^54^, we assessed micro-vessel density (MVD) level by immunohistochemical (IHC) staining of CD34, a molecule that was up-regulated in tumor enriched ECs Endo_c02 and Endo_c03 **(Figure 7c)**, which marked micro-vessels **(Figure 7i)**. The tumor stroma displayed increased vascularization compared to para-cancerous nasal mucosa. Although no statistically significant difference was observed in MVD levels across 3 subtypes (Neural/Basal/Mesenchymal) of ONB patients **(Figure S7c)**, a significant drop of MVD level was found in Hyams IV ONB patients (Hyams I-II vs Hyams IV, *p*=0.026), providing clues of latent ischemic effect for possible treatment resistance and meanwhile, suggesting a potential vulnerability of some well-differentiated tumors to anti-angiogenic therapies. **(Figure 7j)**

### Characterization of Cell-cell Interactions Involved in ONB

We performed a cell-cell interaction analysis so as to explore the interplay among cell types within the tumor microenvironment, which uncovered complicated cell-talk networks between malignant compartment, immune cells and stromal cells in ONB **(Figure 8a)**. When compared with Basal and Neural tumor cells, Mesenchymal malignant cells showed prominent interactions with almost all kinds of TME components, especially with stromal cells including ECs, fibroblasts and SMCs. Moreover, myeloid cells were found to display a central role in cellular communications, with macrophages, monocytes and DCs showing modest number of interactions with all other cell types **(Figure 8b-8c)**.

Thus, we explored the role of tumor and TME cells in remodeling the ecosystem of ONB, by analysis of interaction pairs between tumor and myeloid, stromal cells, respectively **(Figure 8d)**. Cell-cell interactions standing for interplay of oncogenic pathways, including fibroblast growth factors (FGFs), TGF-β receptor, WNT, PDGF and NOTCH signaling pathways were identified. Of note, we found active molecular interactions related to angiogenesis, including VEGFs and MDK (encoding VEGFR) within the network, especially between ECs and malignant cells. Although tumor cells showed strong signals of VEGF-related interactions with multiple TME components regardless of subtypes, Basal and Mesenchymal tumor cells exhibited higher levels of MDK-LRP1 interaction when compared with Neural cells. More up-regulated molecular pairs regarding NOTCH signaling pathway were observed among Basal tumor cells and stromal cells.

Analysis of cellular interaction regarding T cell functions was performed by evaluating expression levels of molecule pairs related to chemokines, T cell co-stimulatory and inhibitory functions **(Figure 8e-8f)**. Instead of cancer cells, myeloid cells especially macrophages were found to occupied the main position to regulate T cell activation an inhibition. For instance, co-stimulatory molecular pairs including TNFSF14-TNFRSF14, ADGRE5-CD55, TNFSF13B-CD40, as well as inhibitory interactions including TNFRSF14-BTLA, TNF-FAS, LGALS9-HAVCR2 and TIGIT-NECTIN2 were detected between CD4&CD8 T cells and myeloid cells, rather than malignant cell. Notably, we found high expression of FAM3C-PDCD1 and low levels of CD274(PD-L1)-PDCD1, PDCD1LG2-PDCD1 between T cells and tumor cells, probably due to the low expression of PD1/PD-L1 on the transcriptomic level^55^. We found exclusively high expression level of inhibitory molecule pair TNF-FAS, and low level of co-stimulatory interaction CD55-ADGRE5, between Basal malignant cells and immune cells, suggestion complex immuno-regulatory status in diverse malignant subtypes.

Overall, cell-cell interaction analysis between malignant cells and major subgroups of TME provided a comprehensive view of ONB ecosystem activity, including tumoral inflammatory response, regulation of T cell function and angiogenesis, from which might derive novel drug targets for ONB. Moreover, Subtle heterogeneity of intercellular interaction among Neural/Basal/Mesenchymal malignant cells has shed light on the importance of biomarkers for more precise prediction of drug efficacy.

## DISSCUSSION

The hypothesis that ONB originated from olfactory epithelium has been widely accepted^56^, which rationalized the comparison between ONB malignant cells and olfactory epithelium components. Clinically, ONB patients presented distinctive sensitivity to chemotherapy and ended with heterogeneous oncologic outcome, indicating the existence of latent tumor subtypes^57^. The prognostic benefit of chemotherapy remained undetermined^57^, and no standard regimen has been recommended. Studies on molecular mechanisms underlying inter- and intra-tumoral differences were constrained by low incidence rate of ONB until the emerging of single-cell RNA sequencing technology, enabling the discrimination of one cellular cluster from others. The scRNA profiles of adult olfactory mucosa epithelium provided by Durante et. al^11^ were extremely suitable to be adopted as a frame of reference for ONB research.

Classe et. al^9^ have reported different clinicopathologic features between two types of ONB, namely Basal and Neural, through a multi-Omics study including mRNA, WES and methylation sequencing, which has set up an innovative classification of ONB phenotypes at bulk resolution. However, we perceived the existence of certain uncategorized samples in their bipartite classification. Hence, scRNA profiles of ONB tumors from a cohort for prospective clinical trial were obtained for further analysis. In the present scRNA study, Neural and Basal signatures proposed in this multi-omics study were found unable to identify all malignant ONB cells, leaving a few tumor subgroups undefined. These previously undescribed cells were found with features of mesenchyme and then matched to olfactory HBC/sustentacular cells by pathway enrichment and correlation analysis. The present study established a novel three-classification of ONB patients, which comprised of Neural, Basal and Mesenchymal subtypes, validated by the external cohort provide by Classe et. al^9^.

In the present study, semi-supervised clustering identified five tumors harboring Mesenchymal gene signatures in bulk ONB samples. CHG and Ki-67 used in the previously mentioned multi-omics research were inherited as important biomarkers for Neural and Basal ONB. IHC staining verified that Mesenchymal ONB samples exhibited abundant CTGF (encoded by CCN2) while little CHG and Ki-67 expression, supporting CCN2 as a representative marker of Mesenchymal ONB. CCN2 served as a multifunctional signaling modulator that can regulate cell proliferation, migration, invasion, drug resistance, and epithelial-mesenchymal transition (EMT)^58^. These three biomarkers better featured ONB phenotypes, as well as assisted in formulating subdivided treatment strategies in clinical practice.

In the present study, malignant ONB cells of Mesenchymal subtype bore some characteristics resembling the sustentacular lineages of OM, including HBCs and two sustentacular clusters. As is well known, HBC contributed significantly in retaining the regenerative process of OM^59^, and upregulation of NOTCH pathway played an essential part in stemness maintenance^59, 60^. We found active NOTCH pathway expression in Mesenchymal malignant subgroups, which might be inherited from HBC, thus leaving a clue on the potential therapeutic target. Besides, some Mesenchymal ONB cells presented increased pro-angiogenetic activity and frequent interactions with endothelial cells, indicating that Mesenchymal cells may take part in tumoral angiogenesis. Higher MVD and lower Hyams grading were detected in Mesenchymal subtype samples than in Basal type. Tentative administration of anti-vessel regimens including endostatin or sunitinib in ONB patients had been occasionally reported^61, 62^, while no survival information was accessible yet. The present study provided theoretical basis of selective usage of anti-angiogenetic drugs in Mesenchymal rather than Basal ONB patients. Therefore, high quality clinical trials were in need.

Neural malignant cells were found with remarkably higher HES6 expression when compared with ORNs. In normal olfactory epithelium, HES6 promoted neuronal differentiation and was an important component of the NOTCH pathway^63^. Although Neural malignant cells perpetuated many features of neural lineages in OM with increase expression of HES6, they still exhibited negative expression of OMP, a marker of mature olfactory neurons, suggesting close relationship between Neural subtype cells and olfactory neurons, while malignant cells might be hindered from becoming mature during tumorigenesis. Increased expression of HES6 has been detected in various tumors including advanced astrocytoma, glioblastoma and prostate cancer, compounding prognostic outcomes via regulating the growth and motile ability of tumor cells^63^. HES6-related activation of NOTCH signaling pathway might represent a latent research avenue toward uncovering therapeutic targets in ONB.

This study also illustrated the differences between normal OM cells and ONB malignant cells, so as to explore commonly expressed signatures and targets shared by all three subtypes. ONB expressed lower epithelial biomarkers like KRTs or EPCAM, and higher interstitial biomarkers like VIM, especially in Mesenchymal subtype tumors, suggesting that ONB underwent an EMT process^10^.

TUBB3 (also known as βIII-tubulin, encoding TUJ1) was overexpressed in all three subtypes of ONB cells when respectively compared with their corresponding putative normal OM origins, in line with a previous research^64^. Generally, in cells of neuronal origin, TUBB3 is reckoned a microtubule protein involved in neurogenesis, axon guidance and maintenance^65^. TUBB3 also played as a biomarker for resistance to microtubule-targeting anti-cancer drugs including vincristine^64^. Several studies have revealed the involvement of TUBB3 over-expression in tumorigenesis and that cancer progression could be reported mitigated by targeting TUBB3-related signaling axis^66^. A study exploring anti-cancer nanomedicines in non-small cell lung cancer has showed that delivery Tubb3 siRNA to lung cancer cells triggered synergistic effect with chemotherapy^67^. The universally high expression of TUBB3 in ONB drew attention to its relationship with clinical outcomes and drug resistance of ONB. Considering that high expression of TUBB3 were related to microtubule-targeting drug resistance, adding vincristine to conventional chemotherapy regimens became of dubious benefit to most ONB patients. Taking the opposite case, innovative targeting therapies including Tubb3 siRNA might play promising roles in the adjuvant treatment of ONB.

TME components in ONB remains far from well recognized, as well as in other sinonasal small round cell tumors. For the first time, this scRNA research unveiled the detailed constitution of T/NK cells and myeloid cells at single-cell resolution. Reported by Classe et al.^9^, most ONB is lack of tumor infiltrating lymphocytes at intraepithelial regions, while several high-grade ONB was highly infiltrated by TILs^68^. Similarly, the present study detected very few TILs, including a small amount of FOXP3-resting and CTLA4-immunosuppressive CD4+ Tregs and relatively abundant immunosuppressive myeloid cells in intraepithelial regions, suggesting that evaluating the abundance of TILs and myeloid cells might be used as indicators for presuming drug responses to different immunotherapeutic strategies.

Notably, a tendency toward increased T cell exhaustion was observed in Tex cells in Basal subtype ONB tumors, when compared to Tex cells in the other two ONB subtypes or the normal OM. Meanwhile, Basal and Neural ONB subtypes harbored higher Treg scores than the Mesenchymal subtype, thus indicating possibility that the ONB patients with basal subtype tumors might benefit most from the therapy combining Treg-cell targeting and effector T cells activation. Tex cells exhibited significantly higher expression levels of LAG3 and PDCD1(encoding PD-1), which were more explicit than TIGIT or CTLA4, implying that immunotherapies targeting LAG3 and PDCD1 might serve as candidates for ONB immune treatment.

The M1 and M2 binary classification of macrophages represented distinctive functional diversions on the spectrum of macrophage polarization^69^. The M2 macrophage phenotype typically produced anti-inflammatory cytokines and played a significant role of immunosuppression. A study focusing on murine protumor macrophages demonstrated that macrophagic S100A4 enhanced M2-like macrophage polarization^42^. Moreover, the overexpression of S100A4 in immunosuppressive macrophages was identified as a regulator of immunosuppressive myeloids in glioblastoma ^70^. In our study, increased expression of S100A4 was observed in TAMs, especially in the Macro-SPP1 subgroup. The Macro-SPP1 subset was found to produce notably high angiogenic factors and exhibited active M2 polarization with very limited phagocytosis characterization. S100A4 was positively correlated with M2 polarization, rendering that macrophagic S100A4 could be a potential therapeutic target to reverse the M2-polarization of TAMs or induce a shift from immunosuppressive TAMs toward anti-tumoral M1-type macrophages in ONB.

Several limitations of the present study should be acknowledged. Firstly, although ONB was regarded genetically resemble to the OM, the origin of the tumor was still not proved yet because of the deficit of experimental models of ONB. Also, whether the three subtypes shared a common cell-of-origin from the GBCs-like tumor cells, or if they arouse separately from different cellular components in the normal OM, are still difficult to be determined. Hence, in vitro and in vivo experiments should be designed for further illumination on this unique malignant tumor.

## CONCLUSION

This study not only established an innovative three-classification system via single-cell analysis and elucidated the markers for categorizing bulk ONB samples into new subtypes with fine feasibility. The projection from scRNA profiles to bulk RNA seq samples enabled the discrimination of tumor phenotypes according to solely IHC results rather than scRNA sequencing. Promisingly, different responses towards certain chemotherapy drugs could be inferred according to the molecular features of three tumor types. Relative deficiency of TILs and abundance of immunosuppressive TAMs indicated the benefits of immunotherapies targeting myeloid cells. However, more evidences including in vitro and in vivo experiments as well as high quality clinical trials were in need before the innovative classification theory could guide accurate treatment of ONB.

## MATERIAL & METHODS

### Patient data & specimens

The study was conducted under the approval of ethics committee of the Eye, Ear, Nose and Throat (EENT) Hospital of Fudan University, Shanghai, China (No.2021038). Samples were retrieved from patients who were pathologically diagnosed with olfactory neuroblastoma (ONB) and received endoscopic resection at the Department of Rhinology of the EENT Hospital of Fudan University with informed consent independently signed and agreed to donate their specimens for scientific study.

Fresh specimens of 10 ONB patients in a prospective clinical study(ChiCTR2100050605) were collected for single-cell RNA sequencing (scRNA) during surgical resection, from July, 2020 to August, 2021, under the supervision of a qualified pathologist. Tumor specimens for bulk RNA-seq analysis were harvested from 23 cases of ONB patients from February, 2019 to January, 2022. Formalin-fixed and paraffin embedded (FFPE) specimens of tumor samples and cancer adjacent mucosa were collected from 30 ONB patients, including 10 patients for scRNA-seq, under the permission of the Pathology Department of the EENT Hospital of Fudan University from February, 2019 to March, 2022, for hematoxylin and eosin (HE) and immunohistochemistry IHC staining. All FFPE specimens were independently reviewed by two pathologists (Y.C.C and C.X.K) who confirmed the diagnosis of ONB and determined relevant Hyams grading of each slide.

Overall, a total of 35 patients with confirmed diagnosis of ONB were enrolled in this study. The collection and utility of patient specimens strictly followed procedures in accordance with the ethical standards formulated in the Helsinki Declaration. The available clinical characteristics of the patients including age, gender, primary tumor localization, orbital invasion, clinical stage including Kadish, Dulguerov and AJCC 8^th^ stage, as well as Hyams grade were summarized in **Supplementary Table S1.**

### RNA Extraction, Bulk RNA-seq and Data Analysis

Total RNA was extracted from ONB tumor tissue samples using RNAiso Plus reagent (Takara Bio, Inc.) according to the manufacturer’s protocol. Sequencing libraries were generated using mRNA-seq Lib Prep Kit for Illumina (ABclonal, USA, Cat. RK20302) following manufacturer’s recommendations and index codes were added to attribute sequences to each sample. The libraries were sequenced on a novaseq 6000 sequencing platform (Illumina, Inc.) using the novaseq 6000 Reagent Kits (Illumina, Inc.) according to the manufacturer’s guidelines. Base calling was performed with Real Time Analysis 3 (RTA3, https://support.illumina.com/downloads.html), and the fastq files were generated by bcl2fastq2 v2.20 (Illumina, Inc., https://support.illumina.com/downloads.html). Alignment of the trimmed RNA-seq reads to the ensembl human genome assembly (GRCh38.p13; https://www.ensembl.org/Homo_sapiens/Info/Index) employed HISAT2 v2.2.1 with the options of ’-k 3 -p 20 --pen-noncansplice 1000000’. The counts of the reads mapped to individual genes were calculated by featureCounts v2.0.3. DEGs were identified by the DESeq2 (version3.15 https://bioconductor.org/packages/DESeq2/), a cut-off of padj <0.05 and |log2(fold change) | >1.5 was applied. FeatureCounts v1.5.0-p3 was used to count the reads numbers mapped to each gene. The expression levels of genes were quantified as fragments per kilobase of exon model per million mapped fragments (FPKM).

### Open-Source Data Acquisition

The scRNA-seq count matrix of one normal adult olfactory mucosa (OM) sample, which has been reported by Durante et al.^11^ in their study and has been deposited to Gene Expression Omnibus database under the accession code GSE139522, was downloaded and used as the control data for scRNA analysis in our study. Only the scRNA-seq which labeled OM2 was re-analyzed as the control dataset. The expression matrix for bulk RNA seq, along with the clinical information of 18 ONB patients reported by Classe et al.^9^, was obtained from GEO dataset GSE118995.

### scRNA Seq: Tissue dissociation and Library Construction

In order to generate single-cell suspensions for single-cell RNA-seq, fresh ONB samples were washed 3 times with phosphate-buffered saline (PBS, Sinopharm, China), and minced into small pieces (< 1 mm^3^) on ice. Samples were then digested mechanically and enzymatically with the Tumor Dissociation Kit, human (Miltenyi Biotec, Germany) following protocols provided by the manufacturer. After digestion, samples were filtered using a 70-mm nylon strainer (Thermo Fisher Scientific, USA), washed with 1% bovine serum albumin (BSA, Sigma-Aldrich, Germany) and 2 mM EDTA in PBS. Optionally, RBCs and debris of dead cells in single cell suspensions were respectively removed using Red Blood Cell Lysis Solution (10x) Kit (Miltenyi Biotec, Germany) and Dead Cell Removal Kit (Miltenyi Biotec, Germany) if needed, according to the manufacturer’s specifications. The cell pellet was resuspended in PBS with 1% BSA and assessed for viability and cell agglomeration rate using a Countess instrument (Thermo Fisher Scientific, USA). Only suspensions with viability ≥ 80% and cell agglomeration rate ≤ 20% were regarded as qualified sample for library construction. Cells of qualified samples were then diluted with PBS containing 1% BSA to about 700∼1200 cells/μl for scRNA sequencing.

Single-cell suspensions were processed using the Chromium Next GEM Single Cell 3’ GEM, Library & Gel Bead Kit v3.1 (10x Genomics, Pleasanton, CA, USA) and run in the Chromium Controller following the manufacturer’s protocol, with a capture target of 8,000-10,000 cells for each sample. A chromium single-cell 3′-library was then constructed with Chromium Next GEM Single Cell 3’ Library Construction Kit v3.1 (10x Genomics, Pleasanton, CA, USA) with the quality-qualified cDNA. After fragmentation, adaptor ligation, sample index PCR, etc., the library is finally quantitatively examined. The final library pool was sequenced on the Illumina Novaseq6000 instrument using 150-base-pair paired-end reads.

### Data processing, Quality Control (QC) and Doublets Filtering

Mapping to homo sapiens GRCh38(v32) genome reference, we performed preliminarily alignment, qualification of the data and read counting of Ensembl genes with CellRanger software (version 5.0.0) using default parameters. Then the raw gene expression matrices of 10 ONB samples and one olfactory mucosa sample (OM2) acquired from GEO database (GSE139522) were merged together and converted to a Seurat object in R (version 3.6.1) using the Seurat package (version 3.2.0)^71, 72^.

Single cells with less than 400 or more than 30,000 UMIs detected, with less than 200 or more than 8,000 genes detected, or with more than 10% mitochondrion derived UMI counts detected have been filtered out as Low-quality cells. Cells exhibiting positive features of both immune and epithelial cells (CD45+, EPCAM+/KRT18+) were screened out as doublets. As a result, a total of 96325 cells including 85452 cells from ONB datasets and 10873 cells from the OM dataset were regarded qualified for downstream analysis. Cells were log normalized to total cellular read-counts and mitochondrial read-counts using the ScaleData() function of the Seurat package.

### Dimensionality Reduction, Clustering and Visualization

Highly variable genes were generated and used for the subsequent principal component analysis (PCA) for dimensionality reduction. The number of appropriate principal components were estimated by an Elbow plot, in combination with exploration of the top genes from each principal component. Clustering was conducted using the FindNeighbors() and FindClusters() functions. The result of clustering was then visualized using Uniform Manifold Approximation and Projection (UMAP) with Seurat functions RunUMAP(). The DimPlot() function was used to generate the visualization format of UMAP plots.

### Cell Type Annotation

Then we performed dimension reduction and re-clustering again for the remaining cells. Differentially expressed genes (DEGs) of a certain cell cluster compared to all other cell clusters were generated using the Seurat function FindAllMarkers. Only DEGs with upregulated expressions were taken account. Major cell types were assigned to Seurat clusters according to the normalized expression of canonical markers, including immune clusters comprised of NK/T cells (PTPRC, CD3D/E/G, NCR1, NCAM1), monocytes (PTPRC, CD14, LYZ, FCN1, S100A8/9), macrophages (PTPRC, CD14, CD68, C1QA/B/C, APOE), neutrophils (PTPRC, CSF3R, S100A8/9), B cells (PTPRC, CD79A/B, MS4A1) and plasma cells (PTPRC, MZB1, CD79A, JCHAIN), stromal clusters consisting of fibroblasts (DCN, COL1A1, PDGFRA), endothelial cells (PECAM1, VWF, CDH5) and smooth muscle cells (RGS5, MYH11, ACTA2). We also found a cluster of olfactory epithelial-like (OE-like) cells (EPCAM, KRT18, KRT8) consisting of blending cell components, with highly expressed DEGs resembling olfactory neural lineages (SOX2, HES6, TUBB2B, NEUROD1/G1, CHGA/B), olfactory sustentacular cells (AQP3, F3, CXCL17, CYP2J2) and Bowman’s glands (CEBPD, NCOA7), according to curated gene sets provided by Durante et al. and Fletcher et al. in their scRNA studies focusing on human and murine olfactory neurogenesis^11, 12^. Before downstream analysis, we filtered putative doublets again following an iterative process introduced by Wu et al^55^. Within each major cell type cluster, we removed the cells with double positive expressions of the canonical marker genes of 11 major cell types discussed above.

### Inferring CNV Based on scRNA-seq Data

Epithelial cells (n = 30660) were selected using Seurat function subset(). The R package inferCNV (version 1.2.1) was adopted to estimate initial CNVs of epithelial cells^15^. In turn, for each ONB sample the epithelial cells were imported as input, together with 500 fibroblasts from OM2 dataset as spike-in normal cells and 800 fibroblasts as normal control. Each single cell was estimated with a CNV score according to the measures introduced by Neftel et al^73^. CNV signal reflecting the overall extent of variations and CNV correlation between each single CNV profile and the average level of all epithelial cells were calculated. CNV observations were sorted by unsupervised clustering based on CNV scores and plotted on a dendrogram. The dendrogram was cut at an appropriate point (k) which was determined through evaluating the similarity between epithelial clusters and spike-in normal cells. Epithelial cells from tumor scRNA profiles that was mixed with normal cells in unsupervised clustering were identified as “non-malignant”, whereas the remaining cells were labeled as “malignant”, thereby annotating the malignancy of each single epithelial cell^15^. Malignant epithelial cells were then re-clustered for downstream analysis.

### Cell Subtype Annotation within Major Cell Clusters

The stromal and immune cell groups identified above, including T cells, myeloid cells, fibroblasts, endothelial cells were further re-clustered to reveal subtypes within major cell clusters. By repeating the abovementioned steps (normalization, dimensionality reduction, and clustering), we further identified and annotated different specific cell subtypes, according to the average expression of canonical marker genes. For each subcluster of a major cell type (e.g. C1 of CD8+ T cells), we assigned a cluster identifier with a marker gene (e.g. GZMK) and its putative functional identity (e.g. effector memory T cells) as “CD8_GZMK_Tem”. The selection criteria of the marker gene for annotation included (1) with top ranking at the differential gene expression analysis for the corresponding cell cluster, (2) with strong specificity of gene expression meaning high expression ratio within the corresponding cell cluster but low in other clusters, and (3) with literature supports that it’s either a marker gene or functional relevant to the type of cell. Markers used in this pipeline were addressed in **Supplementary Table S12**.

### Expression Programs of Intra-tumoral Heterogeneity

Non-negative matrix factorization (NMF) analysis was performed to extract transcriptional programs of malignant cells of each tumor using R package NMF^17, 18^. K = 4 was selected as the number of factors for each tumor, because it yielded a high cophenetic correlation coefficient^74^. A total of 40 metagenes were identified across 10 ONB tumors, and we listed the top-ranked genes according to their loadings of the NMF factor, as shown in **Supplementary Table S5**. Hierarchical clustering of the scores for each program using one minus the Pearson correlation coefficients as the distance metric and Ward’s linkage revealed 5 correlated sets of meta-programs. The gene list of 5 meta-programs can be found in **Supplementary Table S6**.

### Defining functional module scores

The degree to which individual cells express a certain predefined expression program was evaluated using the AddModuleScore() function in Seurat^17, 75^. All analyzed genes were binned based on averaged expression, and the control features were randomly selected from each bin, as described previously^76^. The functional modules including cell cycling score for malignant cells and normal olfactory epithelial cells, cytotoxicity and exhaustion scores for CD8^+^ T cells, Treg and naiveness scores for CD4^+^ T cells, M1, M2, phagocytosis and angiogenesis scores for macrophages, as well as maturation, activation, migration, and chemokine scores for dendritic cells. The involved genes were respectively listed in figures, figure legends or supplementary tables in our manuscript.

### Hybrid Expression Patterns of Neural/Basal/Mesenchymal (NBM) Features in ONB Malignant Cells

The Neural/Basal/Mesenchymal (NBM) Scores were calculated for each single malignant ONB cell, using the Top 50 DEGs among Neural, Basal and Mesenchymal malignant subgroups as gene modules for the Seurat function AddModuleScore(), respectively. Then, the distribution of NBM signatures in each single malignant cell were projected and displayed in a ternary diagram, with each dot (x,y) in the plots representing:

x = Basal Score – Mesenchymal Score;

y = max (Basal Score - Mesenchymal Score) - Neural Score.

The ternary diagram was split to show the hybrid expression patterns of NBM signatures in each ONB samples. Results were visualized mainly using R package ggplot2.

### Neural/Basal/Mesenchymal (NBM) scores defined for bulk ONB samples

Each expression matrix of bulk ONB samples was deemed equivalent to the expression matrx of a single cell, which was then processed using the R package Seurat. Thus, the Seurat function AddModuleScore() were adopted to evaluate the expression levels regarding Neural/Basal/Mesenchymal (NBM) signatures in bulk ONB samples, as described above for single cells.

### Projection of Neural/Basal/Mesenchymal (NBM) Signatures Stratified Bulk ONB samples

Neural/Basal/Mesenchymal (NBM) Signatures defined by scRNA analysis were used to assign ONB molecular subtypes (Neural, Basal and Mesenchymal) to bulk ONB cohort. For our analyses, Top 50 DEGs among NBM subgroups of malignant cells were selected to generate a list of 150 genes. A bulk RNA seq expression matrix of ONB was built according to this NBM signature gene list for semi-supervised hierarchical clustering analysis. The heatmap displaying k-means clustering results were generated using R package ComplexHeatmap, integrating the sample clinical information.

The same hierarchical clustering analysis was also carried out in an external validation ONB bulk cohort deposited by Classe et al. (#GSE118995)^9^. The expression matrix was downloaded from GEO, and the R package limma was adopted for data normalization and scaling. Similar stratification in bulk ONB samples regarding expression levels of NBM signatures was observed in the external validation dataset **(Supplementary Figure S3a)**.

### Assignment of ONB Subtypes

The protein-based expression levels of CHG (marker for Neural subtype of ONB), Ki-67 (marker for Basal subtype of ONB) and CCN2/CTGF (marker for Mesenchymal subtype of ONB) were quantified using Immunohistochemistry (IHC) staining (see later in the method for IHC). The Tumors were first categorized by high Ki-67 level (Basal subtype if Ki-67%≥25%)^9^, hereafter by universally positive expression of Neural biomarker CHG (Neural subtype if H-score of CHG≥75), and then determined by their solely high expression of CCN2 (Mesenchymal subtype if Ki-67%<25%, H-score of CHG<75, H-score of CCN2≥75) **(Supplementary Figure S3b)**.

### Identification of Olfactory Neural and Sustentacular Lineages

In order to distinguish the neural and sustentacular subpopulations in normal olfactory epithelial which have been previously described by Durante et al in their published work^11^, we basically followed the analysis procedures and consulted the parameters provided by the authors in the study. As a result, there was a total of 11348 cells from OM2 dataset remained. By consulting the representative markers of olfactory epithelial cell subpopulations provided by Durante et al^11^, we identified immature (GNG8) and mature olfactory neural lineage cells (GNG13), immediate neuronal precursors (INPs, markers including LHX2 and SSTR2), GBCs (HES6, NEUROD1, NEUROG1, CXCR4), olfactory HBCs (AQP3, F3, SOX2, TP63, CXCL14, KRT5) and sustentacular cells (HES1 and CXCL17). Then we conducted sub-clustering to subdivide these olfactory epithelial cells by referring to multiple known murine marker genes for HBCs, GBCs, INPs as well as iORNs and mORNs. Specific genes for cell-type identification are displayed in **Figure 4b**.

### Construction of Pseudo-time Trajectory

The R package Monocle2 (version 2.14.0) was adopted to perform pseudo-time analysis in order to reveal putative trajectories of differentiation and cell-state transition. The developmental trajectory of CD8+ T cells, CD4+ T cells, macrophages and DCs, were constructed respectively. Practically, the data of the indicated clusters calculated in Seurat was fed directly into Monocle2. The function DDRTree was adopted for dimensionality reduction, after which the cells were placed on pseudo-time trajectories by calling the function orderCells. Then the immune cell differentiation trajectories were visualized by the plot_cell_trajectory function with the default parameters of Monocle2. The pseudo-heatmaps reflecting the expression of marker genes varying along the developmental trajectory were generated by the plot_genes_branched_heatmap function. DEGs over the pseudo-time trajectory between cells in different differentiation fate branches were generated by applying the differentialGeneTest function in Monocle2.

We also applied Slingshot (Version 1.5.1), a cell lineage inference algorithm, to uncover the differentiation trajectories and bifurcations of olfactory neural and sustentacular lineages^77^. Slingshot generated principal curves and cell developmental distances for each lineage, which was then mapped to UMAP projection for visualization. The ordering provided by Slingshot, analogous to pseudotime, is referred to herein as developmental order. Based on the inferred developmental distance, heatmaps displaying the average scaled expression profiles of NBM (Neural/Basal/Mesenchymal) ONB subtypes and cell cycle-related features were produced, with cells ordered according to their developmental positions within the olfactory neuronal and sustentacular cell lineages.

The scRNA profiles of normal adult olfactory epithelial cells (OM2) and one Basal subtype ONB tumor dataset (ONB-311) were integrated using the Seurat function merge(). To place ONB and OM cells in a pseudotime trajectory, the integration was re-processed using R package Monocle2. Expression files were normalized using the estimateSizeFactors() function, with genes expressed in fewer than 3 cells excluded. Using the estimateDispersions() function to estimate negative binomial over-dispersion for each gene, genes with mean expression value > 0.1 and variance greater than the empirical dispersion (the best fit mean-dispersion trend-line) were selected as ordering genes for pseudotime analysis.

### Pathway Enrichment Analysis

To gain functional and mechanistic insights of cell clusters or gene modules, Gene Ontology (GO) and KEGG Pathway enrichment analyses were performed using the R package clusterProfiler, following standard pipeline with default settings^78^. Gene categories associated with p value < 0.05 and FDR < 0.05 were considered statistically significant. To compare the difference of signaling pathway enrichment between cell groups originated from normal and tumor tissue, we performed gene set enrichment analysis (GSEA) using the function clusterProfiler::GSEA(). To explore the heterogeneous expression of malignant cells, we performed gene set variation analysis (GSVA) using the R package GSVA (version 1.34.0) ^79^. Collections of signatures included in pathway enrichment analyses consisted of the Hallmark and Ontology gene sets from the Molecular Signatures Database (MSigDB), gene sets from the Reactome Pathway Database, and gene signatures from the KEGG Pathway Database.

### Evaluation of Cellular Similarity

To quantify cellular similarities between malignant subgroups and olfactory developing cell lineages, we first merged the cell-by-gene expression matrix of the tumor cell dataset and the normal olfactory dataset by their shared gene set^19^. Then, for each expression profile of malignant subgroups, the inter-profile Pearson correlation was calculated with differential expression profiles of each single cell from normal olfactory cell types. We also evaluated the cellular similarities from a different angle by applying the AddModuleScore() function in Seurat to evaluated the aggregated expression levels of malignant signatures of ONB (in forms of Neural, Basal and Mesenchymal scores) in olfactory neural and sustentacular subclusters, as described previously.

### SCENIC Analysis

The SCENIC analysis was run using the Python package ‘‘pySCENIC’’ (version 0.10.3), a lightning-fast python implementation of the SCENIC pipeline^80^, with default parameters based on the motifs database for RcisTarget and GRNboost. The motifs database of Homo sapiens was downloaded from the website https://pyscenic.readthedocs.io/en/latest/. The input matrix was the normalized expression matrix of cells of interest. Regulon specific scores (RSS) were generated using the python function regulon_specificity_scores().

### RNA Velocity Analysis

Prediction of RNA velocities was performed using velocyto (version 0.17.0) in Python and following the scVelo (version 0.2.2) Python pipeline^81^. Firstly, spliced/unspliced reads were annotated by velocyto.py using the output of CellRanger (version 5.0.0), generating BAM files and accompanying GTF, which were then saved as loom files. The loom files were reloaded to scVelo to generate count matrices for spliced and unspliced reads. Next, the count matrices were size-normalized to the median of total molecules across cells. Then, calculation of RNA velocity values for each gene in each cell and embedding RNA velocity vector to low-dimension space were performed. The destination of a single cell was defined according to the Pearson correlation coefficient value between the velocity vector and cell state difference vectors of the column cell. The velocity-based cell transition matrix were calculated by transition_matrix() function of scVelo.

### Cellular Interaction Analysis

The Python package CellPhoneDB (version 2.0) was used to construct intercellular interaction networks in tumor microenvironment of ONB, mainly following the default settings of the software^16^. The potential interaction strength between two cell subsets was predicted based on the expression of ligand-receptor pairs. Then interactions trimmed based on significant sites with p<0.05. Specially, if the malignant cells express the receptor or the ligand, then the corresponding interaction was respectively defined as incoming or outgoing, as described previously^17^. Next, we analyzed ligand-receptor pairs with biologically relevance, including chemokines, co-stimulatory, and inhibitory T cell functions, as well as interaction pairs related to angiogenesis, NOTCH pathway and WNT pathway, between various cell subtypes. The relative expression levels (z-scores) and adjusted p value of ligand/receptor molecule pairs were displayed in forms of dot plots or interaction heatmaps.

### Hematoxylin and Eosin (H&E) Staining, Immunohistochemistry (IHC) and immunofluorescence (IF)

Immunohistochemical (IHC) procedures as well as Hematoxylin and Eosin (H&E) staining were performed on 3-5 µm sections of formalin-fixed, paraffin-embedded human tumor or normal tissues as described previously^82^. The following antibodies were used for IHC: anti-Ki67 (1:1500, Abcam, #ab15580), anti-CTGF/CCN2 (1:200, Thermofisher, #PA5-109248), anti-Chromogranin (1:50, Gene Tech (Shanghai), #GT211429), anti-Tubulin β3/TUBB3 (1:300, BioLegend, #MMS-435P) and anti-CD34 (1: 2000, Abcam, #ab81289). The Dako REAL EnVision System (K5007, DAKO, Glostrup, Denmark) was used as the biotinylated second antibody and color substrate solution. Microvessels were detected by CD34, and microvessel density (MVD) was calculated according to methodology described in our previous studies^82^. Expression levels of CCN2 and CHG were evaluated semi-quantitatively with histochemistry scores (H-scores), based on the percentage of staining positive cells and staining intensity: H-score = ΣPi(i+1). Here, Pi represented the percentage of the number of positive cells in the total number of cells in a selected field, and i stands for staining intensity evaluated by two qualified pathologists respectively. Proliferation index (Ki-67% index) referred to the percentage of Ki-67 positive cells in the total number of cells in a selected field.

Immunofluorescence (IF) specific for markers including CD8 (1:800, Abcam, #ab23T710), CD4 (1:400, Abcam, #ab133616), CD3 (1:200, Abcam, #ab16669), FOXP3 (1:100, Abcam, #ab215206), LAG3 (1:100, Abcam, #ab209236), CD68 (1:00, Abcam, #ab955), VEGFA (1:100, Abcam, #ab52917) and S100A4 (1:1000, Abcam, #ab197896) was performed. Multiple process including dewaxing, rehydration, block of endogenous peroxidase activity and non-specific antigens, retrieval of heating induced epitope were performed on 3-5 µm paraffin-embedded slides. After incubating with primary antibodies and horseradish peroxidase-conjugated secondary antibody, the slides were incubated with tyramide signal amplification (TSA) Fluorochromes (Runnerbio, Shanghai, China) for 10-30 min at room temperature. After the second or third round of antibody incubation and PBS washing, the slides were mounted using Antifade Mounting Medium with DAPI (Beyotime, P0131, Jiangsu, China).

### Survival Analysis

Patients were stratified into three subtypes namely Basal, Neural and Mesenchymal according to their expression patterns of Ki-67/CHG/CCN2, and the disease-free survival (DFS) outcomes of 3 patient groups were presented by survival plots generated by Kaplan-Meier method using R package Survival and Survminer. P-values <0.05 was considered statistically significant.

### Statistics Analysis

Overall, statistical tools, methods, and threshold for each analysis are explicitly described with the results or detailed in the figure legends or Materials and Methods. Statistical comparisons were performed using GraphPad Prism 9 (GraphPad Software, Inc.) or R program (version 3.6.1). Values and error bars represent the mean ± standard error of the mean. The respective number of replicates (n) is indicated in the figures or in the figure legends. p-Values were determined by an appropriate statistical test such as Fisher exact test, one-way ANOVA and log-rank test, as indicated in the figure legends. Detailed descriptions of statistical tests were specified in the results section and in the Figure legends.

---

**Figure S1. Identification and characteristics of cell types in ONB by scRNA analysis.**

a) Quality measures of scRNA-seq experiments, plots showing median numbers of RNA molecule counts, number of genes, and percentage of mitochondrial genes detected in cells of 10 ONB samples.

b) UMAP visualization of the integration of 10 ONB samples and one normal olfactory mucosa sample, colored by different tissue origins and major cell types with results shown by each patient.

c) Violin plots showing differential expression of canonical markers of major cell types in ONB tumor microenvironment.

d) Chromosomal landscapes of inferred large-scale CNVs distinguishing malignant epithelial cells from non-malignant epithelial cells in each ONB tumor sample. The x- and y-axis stands for chromosomal regions and individual cells, respectively. Inferred chromosomal landscapes of ONB-199 (top), ONB-288 (middle) and ONB-946 (bottom) were displayed as representative results. Amplifications (red) or deletions (blue) were inferred by averaging expression over 100-gene stretches on the indicated chromosomes (bottom panels). The inferred CNV pattern of the control fibroblasts extracted from normal OM (OM2) is also shown (top panels).

**Figure S2. scRNA profiles of malignant ONB cell subclusters.**

a) UMAP feature-plots and violin plots showing differential distribution of Neural/Basal features (derived from previous bulk RNA seq analysis) in different ONB malignant subgroups.

b) Heatmaps showing gene expression programs extracted from representative ONB tumors using NMF analysis (ONB-311 and ONB-599, upper and lower plot, respectively).

c) Panels of violin plots depicting the scores for one of the five NMF meta-program signatures for malignant cells from 10 ONB tumors, each panel (from top to bottom) representing one malignant meta-program.

d) UMAP feature-plot and violin plot showing differential distribution of inferred CNV scores in ONB malignant subgroups.

e) Violin plots showing distributions of CNV scores among different cell types from 10 ONB samples.

**Figure S3. Characterization of Neural/Basal/Mesenchymal subtypes of ONB.**

a) Projection of scRNA-seq NBM malignant features onto an external validation bulk ONB cohort (GSE118995). The upper panel: The clinical characteristics of 18 ONB patients including Hyams grade and Dulguerov stage were indicated as different colors. The lower panel: Heatmap showing expression score of each gene of NBM Top 50 signatures **(Table S8)** of each ONB patient in the validation bulk cohort. Patients were stratified into 3 groups using semi-supervised clustering.

b) A flowchart illustrating how ONB tumors were classified into 3 molecular subtypes (Basal, Neural and Mesenchymal).

**Figure S4. scRNA analysis of similarity and divergence between adult human olfactory neuroepithelium and ONB tumor cells.**

a) UMAP displaying 11,348 adult olfactory mucosal cells; N = 1 sample (patient OM2 from the scRNA dataset (GSE139522). The cell cluster phenotype is noted on the color key legend and labels.

b) UMAP visualization, colors representing the expression levels of canonical markers (gray to purple) olfactory neurons, sustentacular cells and primitive basal cells.

c) UMAP feature-plots and violin plots showing differential distribution of Neural/Basal/Mesenchymal malignant features in human olfactory neuroepithelial compartments, in forms of NBM malignant scores.

d) Representative immunohistochemistry (IHC) images displaying wide expression of TUJ1 (encoded by TUBB3) in ONB tumor cells, showing in 2 typical cases of ONB samples. Scale bars, 100 µm (400x).

e) Representative immunohistochemistry (IHC) images displaying scarce expression of TUJ1 (encoded by TUBB3) in sinonasal mucosa melanoma (SNMM) tumor cells, showing in 2 typical cases. Scale bars, 100 µm (400x).

f) Developmental trajectories of olfactory neuronal cells and sustentacular cells from OM2, and ONB tumor cells from ONB-311. Normal OM and malignant cells were shown separately for each subcluster.

g) Developmental trajectory of olfactory epithelial cells and malignant cells from ONB-311. Color gradient indicated the expression level of features regarding cell proliferation.

**Figure S5. scRNA profiles of T/NK cell subclusters in ONB tumor microenvironment.**

a) Violin plots displaying expression levels of Tem-related signature genes (GZMK, PDCD1, LAG3) in CD8^+^ T cell subclusters.

b) Developmental trajectory of CD8^+^ T cells with color gradient indicating the expression level of signatures related to T cell cytotoxicity (GNLY, GZMB, GZMH, PRF1) and exhaustion (CTLA4, LAG3, PDCD1, TIGIT).

c) Spearman correlation between activity of CD8^+^ T cells, as measured by the average granzyme expression (GZMA, GZMB and GZMH) and the expression of CD8^+^ T-cell specific genes (>3-fold overexpressed versus all other cells). Top 20 correlated genes were highlighted as well as genes encoding known immune checkpoint molecules (LAG3 and TIGIT).

d) Representative images of multiplex immunofluorescence (IF) staining of CD8^+^LAG3^+^ T cells in ONB tumor tissue. Proteins detected using respective antibodies in the assays are indicated on top. Scale bars, 10 µm.

e) Developmental trajectory of CD4^+^ T cells with color gradient indicating the expression level of signatures related to Treg function (BATF, CTLA4, LAYN, TNFRSF9).

f) Scatter plot showing the specificity scores of regulons of CD8-PDCD1-Tex, CD8-ZNF683-Trm, CD8-GZMK-Tem, CD8-GZMH-Teff, CD4-IL21-Tfh, CD4-IL7R-Tcm, CD4-FOXP3-resting Treg, and CD4-CTLA4-suppressive Tregs. The top 10 regulons are highlighted.

g) Violin plots displaying expression levels of Exhaustion and Cytotoxic Scores in CD8^+^ T cell subclusters and Naiveness and Treg Scores in CD4^+^ T cell subclusters in ONB tumor microenvironment.

**Figure S6. scRNA profiles of myeloid subclusters in ONB tumor microenvironment.**

a) UMAP visualization of macrophages, DCs and monocyte subclusters split by patient origins, colored by cell subtypes.

b) Violin plots displaying expression levels of marker genes in macrophage subclusters in ONB tumor microenvironment.

c) Developmental trajectory of macrophages with color gradient indicating the expression level of M2 TAM signature SPP1.

d) Violin plots showing the distribution of S100A4 in macrophages, stratified by normal and tumor tissue origins.

e) Representative images of multiplex immunofluorescence (IF) staining of macrophages in ONB tissues. Proteins detected using respective antibodies in the assays are indicated on top. Scale bars, 10 µm.

f) Boxplots and violin plots comparing M1 & M2 Scores, Phagocytosis & Angiogenesis Scores among normal and ONB derived macrophages. P values were calculated using the Wilcoxon rank-sum test.

g) Heatmap showing the selected signaling pathways that were significantly enriched in GSVA analyses for each DC subcluster derived from Hallmark, KEGG and Reactome signaling pathways. Filled colors from dark-blue to red represent scaled expression levels (normalized −log10 p values) from low to high.

**Figure S7. scRNA profiles of stromal subclusters in ONB tumor microenvironment.**

a) UMAP visualization of EC and fibroblast subclusters split by patient origins, colored by cell types.

b) Violin plots displaying expression levels of marker genes in fibroblast subclusters in ONB tumor microenvironment.

c) Boxplots comparing MVD levels among Basal, Neural and Mesenchymal ONB subtypes. P value was calculated using the one-way ANOVA test.

n.s. no significance.

## Reference

1. Dulguerov, P., Allal, A.S. & Calcaterra, T.C. Esthesioneuroblastoma: a meta-analysis and review. The Lancet. Oncology 2, 683–690 (2001).

2. Jethanamest, D., Morris, L.G., Sikora, A.G. & Kutler, D.I. Esthesioneuroblastoma: a population-based analysis of survival and prognostic factors. Archives of otolaryngology--head & neck surgery 133, 276–280 (2007).

3. Turner, J.H. & Reh, D.D. Incidence and survival in patients with sinonasal cancer: a historical analysis of population-based data. Head & neck 34, 877–885 (2012).

4. Abdelmeguid, A.S. Olfactory Neuroblastoma. Current oncology reports 20, 7 (2018).

5. Wu, K., et al. Orbital invasion by Esthesioneuroblastoma: a comparative case series and review of literature. Orbit (Amsterdam, Netherlands), 1–14 (2020).

6. Colevas, A.D., et al. NCCN Guidelines Insights: Head and Neck Cancers, Version 1.2018. Journal of the National Comprehensive Cancer Network : JNCCN 16, 479–490 (2018).

7. Gallia, G.L., et al. Genomic analysis identifies frequent deletions of Dystrophin in olfactory neuroblastoma. Nature communications 9, 5410 (2018).

8. Zhang, Q., et al. Single-cell transcriptome reveals cellular hierarchies and guides p-EMT-targeted trial in skull base chordoma. Cell discovery 8, 94 (2022).

9. Classe, M., et al. Integrated Multi-omic Analysis of Esthesioneuroblastomas Identifies Two Subgroups Linked to Cell Ontogeny. Cell reports 25, 811–821.e815 (2018).

10. Romani, C., et al. Gene Expression Profiling of Olfactory Neuroblastoma Helps Identify Prognostic Pathways and Define Potentially Therapeutic Targets. Cancers 13(2021).

11. Durante, M.A., et al. Single-cell analysis of olfactory neurogenesis and differentiation in adult humans. Nature Neuroscience 23, 323–326 (2020).

12. Fletcher, R.B., et al. Deconstructing Olfactory Stem Cell Trajectories at Single-Cell Resolution. Cell stem cell 20, 817–830.e818 (2017).

13. Gadye, L., et al. Injury Activates Transient Olfactory Stem Cell States with Diverse Lineage Capacities. Cell stem cell 21, 775–790.e779 (2017).

14. Zhang, X., et al. CellMarker: a manually curated resource of cell markers in human and mouse. Nucleic acids research 47, D721–d728 (2019).

15. Maynard, A., et al. Therapy-Induced Evolution of Human Lung Cancer Revealed by Single-Cell RNA Sequencing. Cell 182, 1232–1251.e1222 (2020).

16. Patel, A.P., et al. Single-cell RNA-seq highlights intratumoral heterogeneity in primary glioblastoma. *Science (New York*, N.Y.) 344, 1396–1401 (2014).

17. Puram, S.V., et al. Single-Cell Transcriptomic Analysis of Primary and Metastatic Tumor Ecosystems in Head and Neck Cancer. Cell 171, 1611–1624.e1624 (2017).

18. Jessa, S., et al. Stalled developmental programs at the root of pediatric brain tumors. Nature Genetics 51, 1702–1713 (2019).

19. Dong, R., et al. Single-Cell Characterization of Malignant Phenotypes and Developmental Trajectories of Adrenal Neuroblastoma. Cancer Cell (2020).

20. Xing, X., et al. Decoding the multicellular ecosystem of lung adenocarcinoma manifested as pulmonary subsolid nodules by single-cell RNA sequencing. Science advances 7(2021).

21. Peng, W.S., et al. Dissecting the heterogeneity of the microenvironment in primary and recurrent nasopharyngeal carcinomas using single-cell RNA sequencing. Oncoimmunology 11, 2026583 (2022).

22. Network., C.G.A.R. Comprehensive molecular characterization of gastric adenocarcinoma. Nature 513, 202–209 (2014).

23. Cheng, S., et al. A pan-cancer single-cell transcriptional atlas of tumor infiltrating myeloid cells. Cell 184, 792–809.e723 (2021).

24. Chen, Y.P., et al. Single-cell transcriptomics reveals regulators underlying immune cell diversity and immune subtypes associated with prognosis in nasopharyngeal carcinoma. Cell research (2020).

25. Guo, X., et al. Global characterization of T cells in non-small-cell lung cancer by single-cell sequencing. Nature Medicine 24, 978–985 (2018).

26. Zhang, L., et al. Lineage tracking reveals dynamic relationships of T cells in colorectal cancer. Nature 564, 268–272 (2018).

27. Zheng, C., et al. Landscape of Infiltrating T Cells in Liver Cancer Revealed by Single-Cell Sequencing. Cell 169, 1342–1356.e1316 (2017).

28. Chen, Z., et al. Dissecting the single-cell transcriptome network underlying esophagus non-malignant tissues and esophageal squamous cell carcinoma. EBioMedicine 69, 103459 (2021).

29. van der Leun, A.M., Thommen, D.S. & Schumacher, T.N. CD8(+) T cell states in human cancer: insights from single-cell analysis. Nature reviews. Cancer 20, 218–232 (2020).

30. Liu, S., Liu, X., Zhang, C., Shan, W. & Qiu, X. T-Cell Exhaustion Status Under High and Low Levels of Hypoxia-Inducible Factor 1α Expression in Glioma. Frontiers in pharmacology 12, 711772 (2021).

31. Li, H., et al. Dysfunctional CD8 T Cells Form a Proliferative, Dynamically Regulated Compartment within Human Melanoma. Cell 176, 775–789.e718 (2019).

32. Savas, P., et al. Single-cell profiling of breast cancer T cells reveals a tissue-resident memory subset associated with improved prognosis. Nature Medicine 24, 986–993 (2018).

33. Lambrechts, D., et al. Phenotype molding of stromal cells in the lung tumor microenvironment. Nature Medicine 24, 1277–1289 (2018).

34. Terawaki, S., et al. IFN-α directly promotes programmed cell death-1 transcription and limits the duration of T cell-mediated immunity. *Journal of immunology (Baltimore*, Md. : 1950) 186, 2772–2779 (2011).

35. Zheng, L., et al. Pan-cancer single-cell landscape of tumor-infiltrating T cells. *Science (New York*, N.Y.) 374, abe6474 (2021).

36. Oestreich, K.J., Yoon, H., Ahmed, R. & Boss, J.M. NFATc1 regulates PD-1 expression upon T cell activation. *Journal of immunology (Baltimore*, Md. : *1950)* 181, 4832–4839 (2008).

37. Pittet, M.J., Michielin, O. & Migliorini, D. Clinical relevance of tumour-associated macrophages. Nature reviews. Clinical oncology (2022).

38. Bassler, K., Schulte-Schrepping, J., Warnat-Herresthal, S., Aschenbrenner, A.C. & Schultze, J.L. The Myeloid Cell Compartment-Cell by Cell. Annual review of immunology 37, 269–293 (2019).

39. Yunna, C., Mengru, H., Lei, W. & Weidong, C. Macrophage M1/M2 polarization. European journal of pharmacology 877, 173090 (2020).

40. Shapouri-Moghaddam, A., et al. Macrophage plasticity, polarization, and function in health and disease. Journal of cellular physiology 233, 6425–6440 (2018).

41. Zhang, L., et al. Single-Cell Analyses Inform Mechanisms of Myeloid-Targeted Therapies in Colon Cancer. Cell 181, 442–459.e429 (2020).

42. Liu, S., et al. S100A4 enhances protumor macrophage polarization by control of PPAR-γ-dependent induction of fatty acid oxidation. Journal for immunotherapy of cancer 9(2021).

43. Pathria, P., Louis, T.L. & Varner, J.A. Targeting Tumor-Associated Macrophages in Cancer. Trends in immunology 40, 310–327 (2019).

44. Martin, P. & Gurevich, D.B. Macrophage regulation of angiogenesis in health and disease. Seminars in cell & developmental biology 119, 101–110 (2021).

45. Gardner, A., de Mingo Pulido, Á. & Ruffell, B. Dendritic Cells and Their Role in Immunotherapy. Frontiers in immunology 11, 924 (2020).

46. Zilionis, R., et al. Single-Cell Transcriptomics of Human and Mouse Lung Cancers Reveals Conserved Myeloid Populations across Individuals and Species. Immunity 50, 1317–1334.e1310 (2019).

47. Liu, Y., et al. Tumour heterogeneity and intercellular networks of nasopharyngeal carcinoma at single cell resolution. Nature communications 12, 741 (2021).

48. Allan, R.S., et al. Migratory dendritic cells transfer antigen to a lymph node-resident dendritic cell population for efficient CTL priming. Immunity 25, 153–162 (2006).

49. Zhao, Q., et al. Single-Cell Transcriptome Analyses Reveal Endothelial Cell Heterogeneity in Tumors and Changes following Antiangiogenic Treatment. Cancer research 78, 2370–2382 (2018).

50. Litviňuková, M., et al. Cells of the adult human heart. Nature 588, 466–472 (2020).

51. Gunaydin, G. CAFs Interacting With TAMs in Tumor Microenvironment to Enhance Tumorigenesis and Immune Evasion. Frontiers in oncology 11, 668349 (2021).

52. Mizoguchi, F., et al. Functionally distinct disease-associated fibroblast subsets in rheumatoid arthritis. Nature communications 9, 789 (2018).

53. Lee, J.J., et al. Elucidation of Tumor-Stromal Heterogeneity and the Ligand-Receptor Interactome by Single-Cell Transcriptomics in Real-world Pancreatic Cancer Biopsies. Clinical cancer research : an official journal of the American Association for Cancer Research (2021).

54. Walia, A., et al. Endostatin’s emerging roles in angiogenesis, lymphangiogenesis, disease, and clinical applications. Biochimica et biophysica acta 1850, 2422–2438 (2015).

55. Wu, F., et al. Single-cell profiling of tumor heterogeneity and the microenvironment in advanced non-small cell lung cancer. Nature communications 12, 2540 (2021).

56. Capper, D., et al. DNA methylation-based reclassification of olfactory neuroblastoma. Acta neuropathologica 136, 255–271 (2018).

57. Su, S.Y., et al. Outcomes for olfactory neuroblastoma treated with induction chemotherapy. Head Neck 39, 1671–1679 (2017).

58. Shen, Y.W., Zhou, Y.D., Chen, H.Z., Luan, X. & Zhang, W.D. Targeting CTGF in Cancer: An Emerging Therapeutic Opportunity. Trends Cancer 7, 511–524 (2021).

59. Iwai, N., Zhou, Z., Roop, D.R. & Behringer, R.R. Horizontal basal cells are multipotent progenitors in normal and injured adult olfactory epithelium. Stem Cells 26, 1298–1306 (2008).

60. Dai, Q., et al. Notch Signaling Regulates Lgr5(+) Olfactory Epithelium Progenitor/Stem Cell Turnover and Mediates Recovery of Lesioned Olfactory Epithelium in Mouse Model. Stem Cells 36, 1259–1272 (2018).

61. Sun, Y., et al. Outcomes and Quality-of-Life Measures after Endoscopic Endonasal Resection of Kadish Stage C Olfactory Neuroblastomas. World Neurosurg 151, e58–e67 (2021).

62. Wang, L., et al. Recurrent Olfactory Neuroblastoma Treated With Cetuximab and Sunitinib: A Case Report. Medicine (Baltimore) 95, e3536 (2016).

63. Krossa, I., et al. Recent advances in understanding the role of HES6 in cancers. Theranostics 12, 4374–4385 (2022).

64. Topcagic, J., et al. Comprehensive molecular profiling of advanced/metastatic olfactory neuroblastomas. PLoS One 13, e0191244 (2018).

65. Lin, B., et al. Injury Induces Endogenous Reprogramming and Dedifferentiation of Neuronal Progenitors to Multipotency. Cell Stem Cell 21, 761–774 e765 (2017).

66. Huang, J., et al. Targeting the IL-1beta/EHD1/TUBB3 axis overcomes resistance to EGFR-TKI in NSCLC. Oncogene 39, 1739–1755 (2020).

67. Conte, C., et al. Multi-component bioresponsive nanoparticles for synchronous delivery of docetaxel and TUBB3 siRNA to lung cancer cells. Nanoscale 13, 11414–11426 (2021).

68. Classe, M., et al. Evaluating the prognostic potential of the Ki67 proliferation index and tumour-infiltrating lymphocytes in olfactory neuroblastoma. Histopathology 75, 853–864 (2019).

69. Kiss, M., Van Gassen, S., Movahedi, K., Saeys, Y. & Laoui, D. Myeloid cell heterogeneity in cancer: not a single cell alike. Cell Immunol 330, 188–201 (2018).

70. Abdelfattah, N., et al. Single-cell analysis of human glioma and immune cells identifies S100A4 as an immunotherapy target. Nat Commun 13, 767 (2022).

71. Stuart, T., et al. Comprehensive Integration of Single-Cell Data. Cell 177, 1888–1902.e1821 (2019).

72. Butler, A., Hoffman, P., Smibert, P., Papalexi, E. & Satija, R. Integrating single-cell transcriptomic data across different conditions, technologies, and species. Nature biotechnology 36, 411–420 (2018).

73. Neftel, C., et al. An Integrative Model of Cellular States, Plasticity, and Genetics for Glioblastoma. Cell 178, 835–849.e821 (2019).

74. Brunet, J.P., Tamayo, P., Golub, T.R. & Mesirov, J.P. Metagenes and molecular pattern discovery using matrix factorization. Proceedings of the National Academy of Sciences of the United States of America 101, 4164–4169 (2004).

75. Hovestadt, V., et al. Resolving medulloblastoma cellular architecture by single-cell genomics. Nature 572, 74–79 (2019).

76. Tirosh, I., et al. Dissecting the multicellular ecosystem of metastatic melanoma by single-cell RNA-seq. *Science (New York*, N.Y.) 352, 189–196 (2016).

77. Street, K., et al. Slingshot: cell lineage and pseudotime inference for single-cell transcriptomics. BMC genomics 19, 477 (2018).

78. Yu, G., Wang, L.G., Han, Y. & He, Q.Y. clusterProfiler: an R package for comparing biological themes among gene clusters. Omics : a journal of integrative biology 16, 284–287 (2012).

79. Hänzelmann, S., Castelo, R. & Guinney, J. GSVA: gene set variation analysis for microarray and RNA-seq data. BMC bioinformatics 14, 7 (2013).

80. Aibar, S., et al. SCENIC: single-cell regulatory network inference and clustering. Nature methods 14, 1083–1086 (2017).

81. Bergen, V., Lange, M., Peidli, S., Wolf, F.A. & Theis, F.J. Generalizing RNA velocity to transient cell states through dynamical modeling. Nature biotechnology 38, 1408–1414 (2020).

82. Song, X., et al. Hypoxia-Inducible Factor-1α (HIF-1α) Expression on Endothelial Cells in Juvenile Nasopharyngeal Angiofibroma: A Review of 70 cases and Tissue Microarray Analysis. The Annals of otology, rhinology, and laryngology 127, 357–366 (2018).

