## Supplementary material for "Single-cell transcriptomic landscape deciphers novel olfactory neuroblastoma subtypes and intratumoral heterogeneity": Figure S1

### Supplementary Figure S1

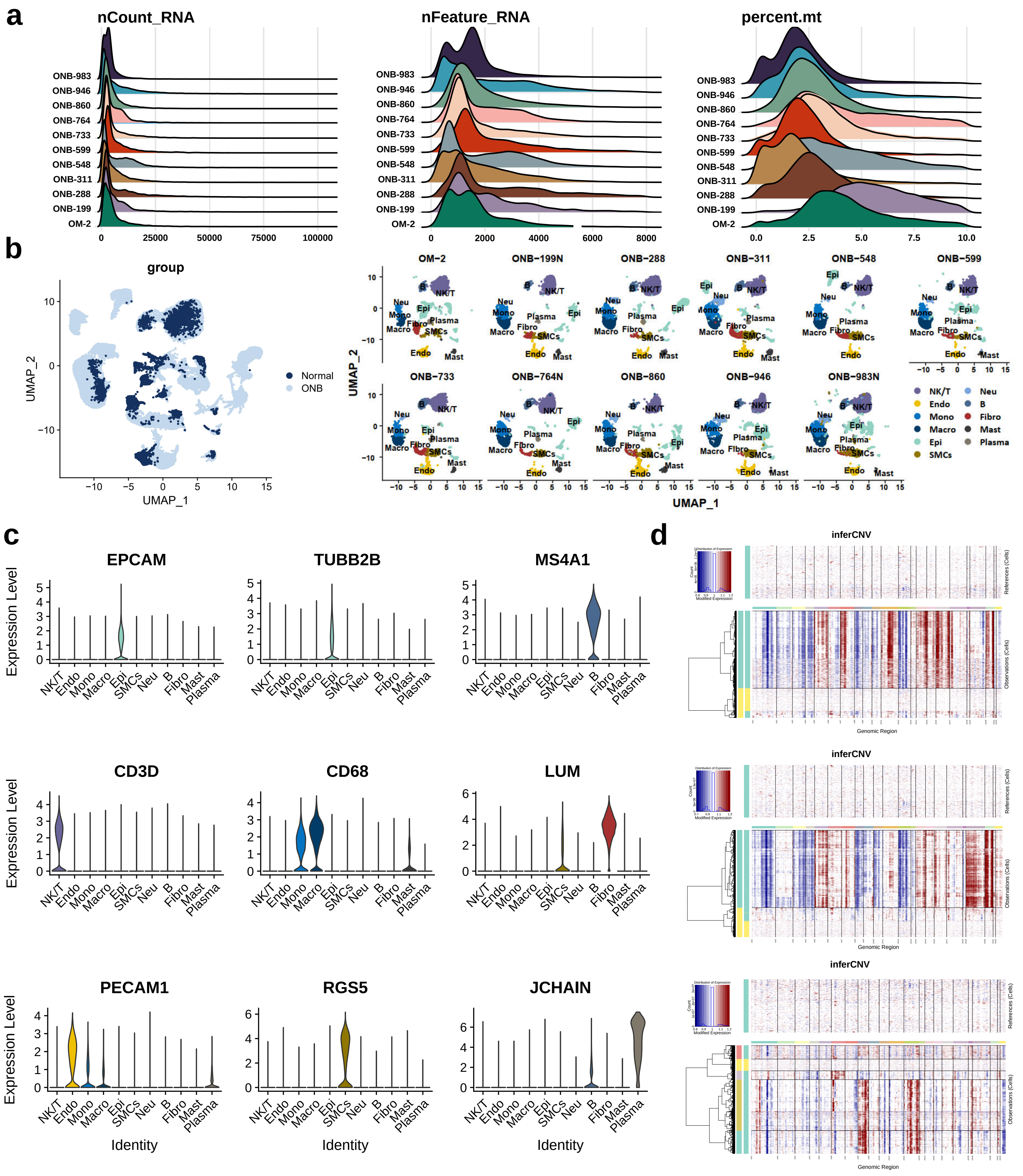

**Supplementary Figure S1.** Identification and characteristics of cell types in ONB by scRNA analysis.

**a)** Quality measures of scRNA-seq experiments, plots showing median numbers of RNA molecule counts, number of genes, and percentage of mitochondrial genes detected in cells of 10 ONB samples.

**b)** UMAP visualization of the integration of 10 ONB samples and one normal olfactory mucosa sample, colored by different tissue origins and major cell types with results shown by each patient.

**c)** Violin plots showing differential expression of canonical markers of major cell types in ONB tumor microenvironment.

**d)** Chromosomal landscapes of inferred large-scale CNVs distinguishing malignant epithelial cells from non-malignant epithelial cells in each ONB tumor sample. The x- and y-axis stands for chromosomal regions and individual cells, respectively. Inferred chromosomal landscapes of ONB-199 (top), ONB-288 (middle) and ONB-946 (bottom) were displayed as representative results. Amplifications (red) or deletions (blue) were inferred by averaging expression over 100-gene stretches on the indicated chromosomes (bottom panels). The inferred CNV pattern of the control fibroblasts extracted from normal OM (OM2) is also shown (top panels).
