## Supplementary material for "Single-cell transcriptomic landscape deciphers novel olfactory neuroblastoma subtypes and intratumoral heterogeneity": Figure S2

### Supplementary Figure S2

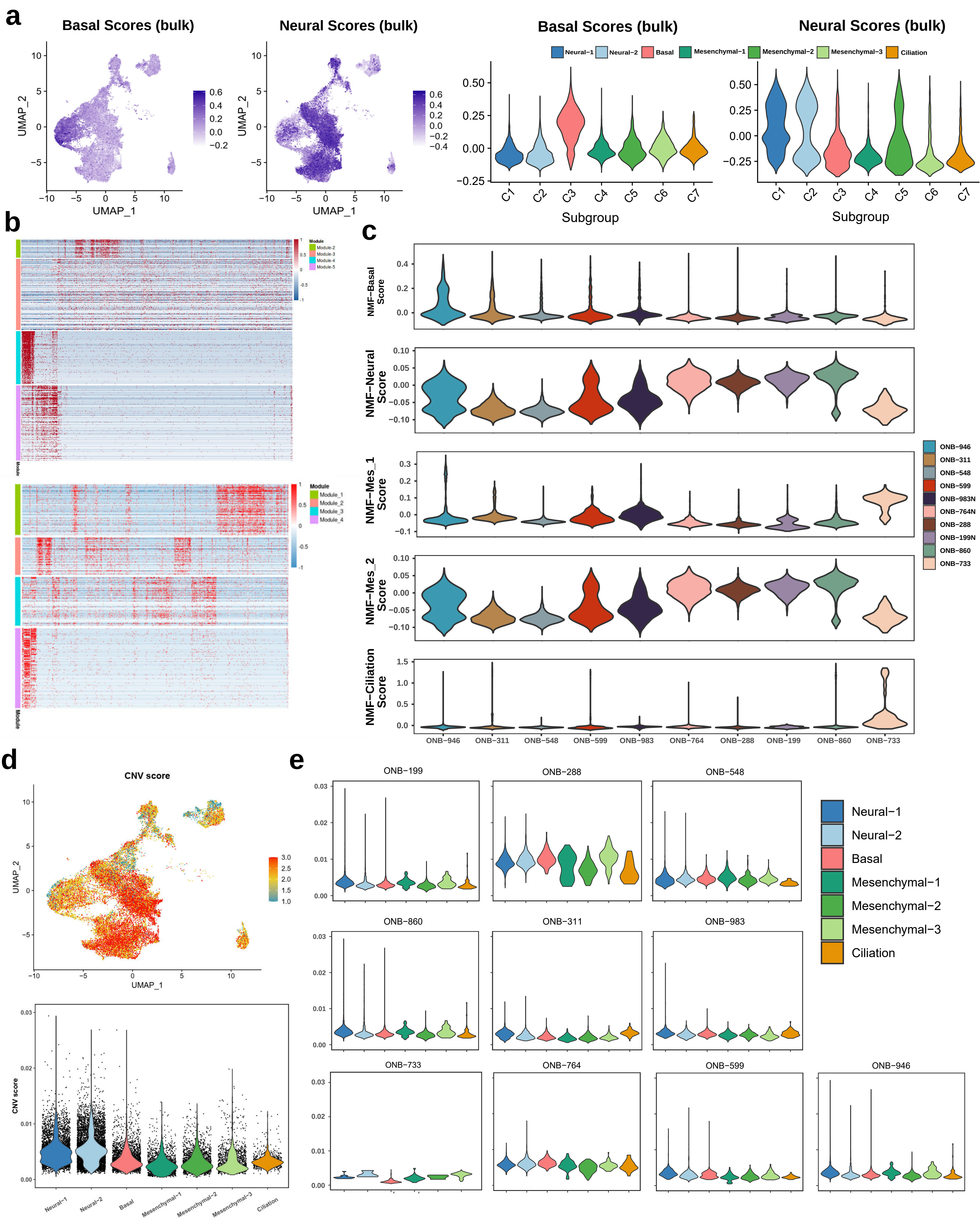

**Supplementary Figure S2.** scRNA profiles of malignant ONB cell subclusters.

**a)** UMAP feature-plots and violin plots showing differential distribution of Neural/Basal features (derived from previous bulk RNA seq analysis) in different ONB malignant subgroups.

**b)** Heatmaps showing gene expression programs extracted from representative ONB tumors using NMF analysis (ONB-311 and ONB-599, upper and lower plot, respectively).

**c)** Panels of violin plots depicting the scores for one of the five NMF meta-program signatures for malignant cells from 10 ONB tumors, each panel (from top to bottom) representing one malignant meta-program.

**d)** UMAP feature-plot and violin plot showing differential distribution of inferred CNV scores in ONB malignant subgroups.

**e)** Violin plots showing distributions of CNV scores among different cell types from 10 ONB samples.
