## Supplementary material for "Single-cell transcriptomic landscape deciphers novel olfactory neuroblastoma subtypes and intratumoral heterogeneity": Figure S3

### Supplementary Figure S3

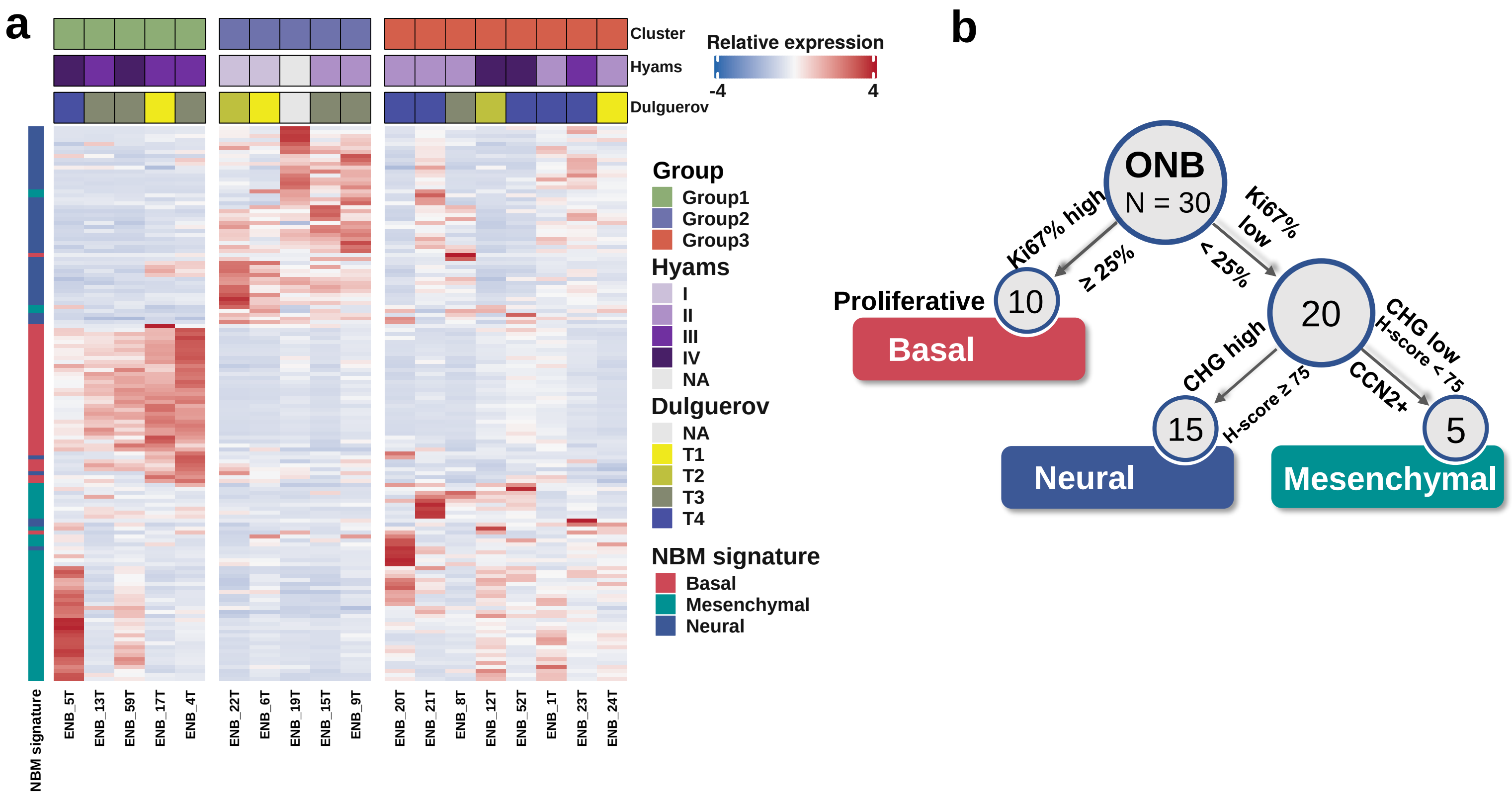


**Supplementary Figure S3.** Characterization of Neural/Basal/Mesenchymal subtypes of ONB.

**a)** Projection of scRNA-seq NBM malignant features onto an external validation bulk ONB cohort (GSE118995). The upper panel: The clinical characteristics of 18 ONB patients including Hyams grade and Dulguerov stage were indicated as different colors. The lower panel: Heatmap showing expression score of each gene of NBM Top 50 signatures **(Table S8)** of each ONB patient in the validation bulk cohort. Patients were stratified into 3 groups using semi-supervised clustering.

**b)** A flowchart illustrating how ONB tumors were classified into 3 molecular subtypes (Basal, Neural and Mesenchymal).
