## Supplementary material for "Single-cell transcriptomic landscape deciphers novel olfactory neuroblastoma subtypes and intratumoral heterogeneity": Figure S4

### Supplementary Figure S4

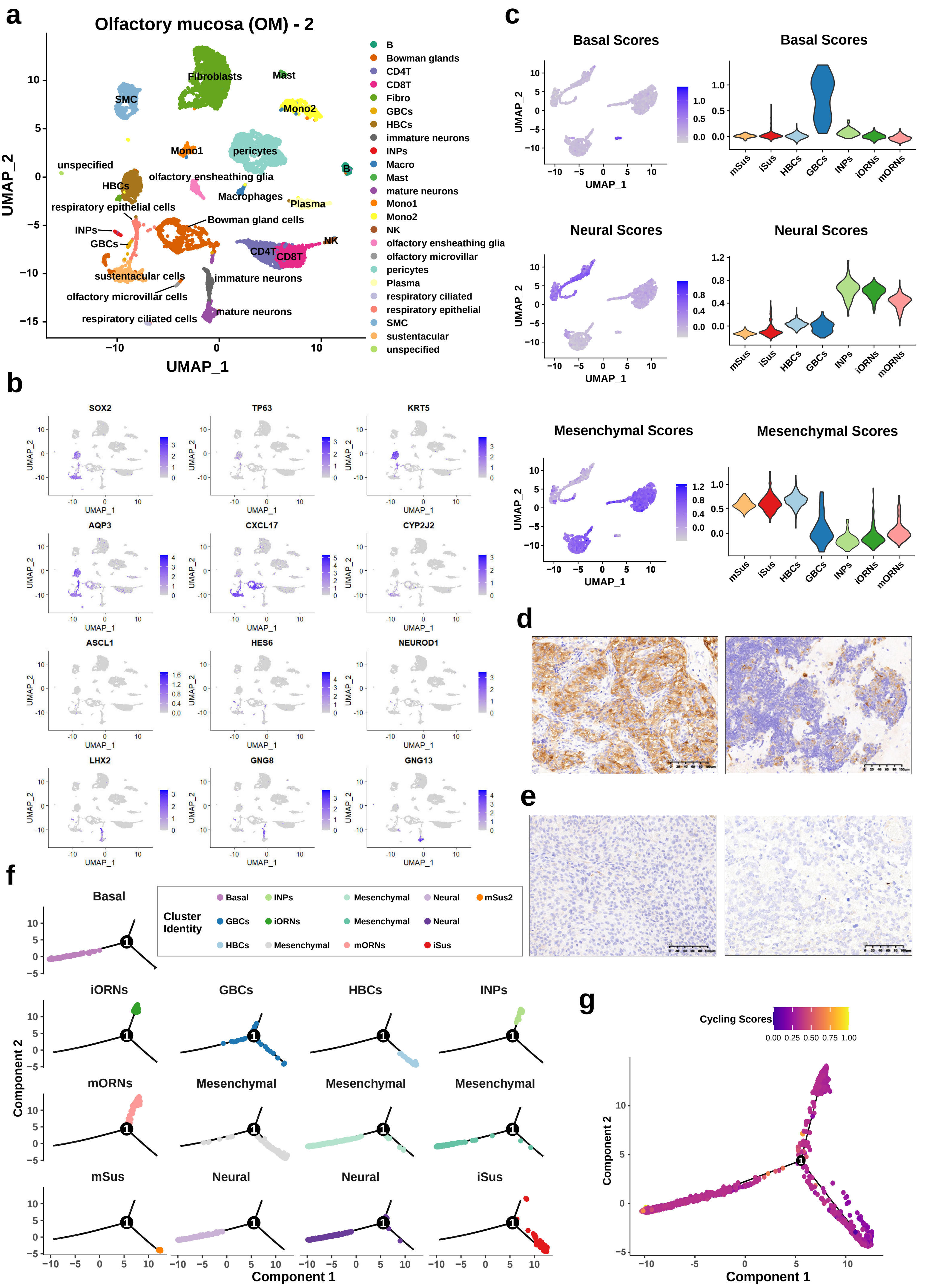

**Supplementary Figure S4.** scRNA analysis of similarity and divergence between adult human olfactory neuroepithelium and ONB tumor cells.

a) UMAP displaying 11,348 adult olfactory mucosal cells; N = 1 sample (patient OM2 from the scRNA dataset (GSE139522). The cell cluster phenotype is noted on the color key legend and labels.

b) UMAP visualization, colors representing the expression levels of canonical markers (gray to purple) olfactory neurons, sustentacular cells and primitive basal cells.

c) UMAP feature-plots and violin plots showing differential distribution of Neural/Basal/Mesenchymal malignant features in human olfactory neuroepithelial compartments, in forms of NBM malignant scores.

d) Representative immunohistochemistry (IHC) images displaying wide expression of TUJ1 (encoded by TUBB3) in ONB tumor cells, showing in 2 typical cases of ONB samples. Scale bars, 100 μm (400x).

e) Representative immunohistochemistry (IHC) images displaying scarce expression of TUJ1 (encoded by TUBB3) in sinonasal mucosa melanoma (SNMM) tumor cells, showing in 2 typical cases. Scale bars, 100 μm (400x).

f) Developmental trajectories of olfactory neuronal cells and sustentacular cells from OM2, and ONB tumor cells from ONB-311. Normal OM and malignant cells were shown separately for each subcluster.

g) Developmental trajectory of olfactory epithelial cells and malignant cells from ONB-311. Color gradient indicated the expression level of features regarding cell proliferation.
