## Supplementary material for "Single-cell transcriptomic landscape deciphers novel olfactory neuroblastoma subtypes and intratumoral heterogeneity": Figure S5

### Supplementary Figure S5

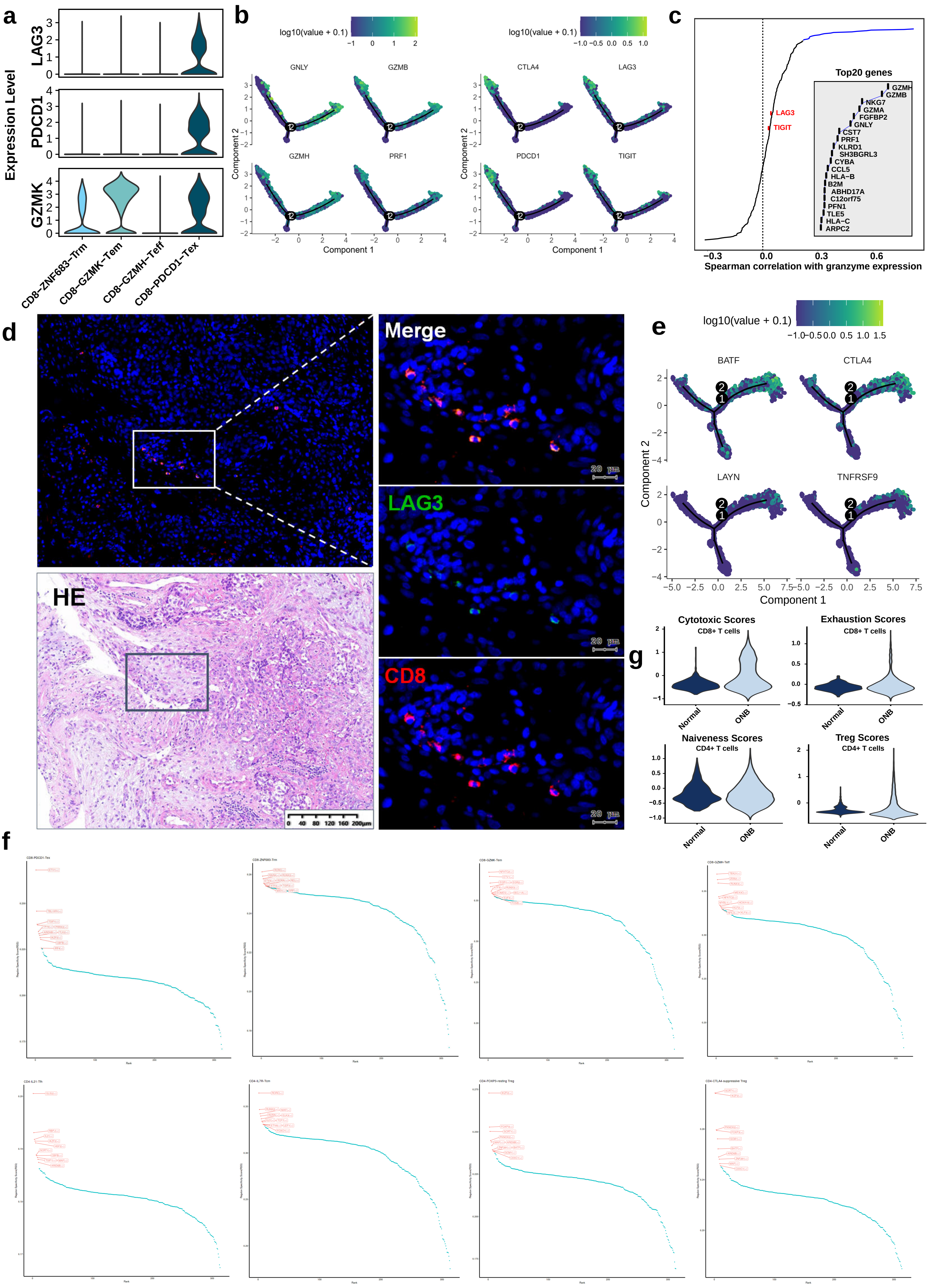

Supplementary Figure S5. scRNA profiles of T/NK cell subclusters in ONB tumor microenvironment.

a) Violin plots displaying expression levels of Tem-related signature genes (GZMK, PDCD1, LAG3) in CD8+ T cell subclusters.

b) Developmental trajectory of CD8+ T cells with color gradient indicating the expression level of signatures related to T cell cytotoxicity (GNLY, GZMB, GZMH, PRF1) and exhaustion (CTLA4, LAG3, PDCD1, TIGIT).

c) Spearman correlation between activity of CD8+ T cells, as measured by the average granzyme expression (GZMA, GZMB and GZMH) and the expression of CD8+ T-cell specific genes (>3-fold overexpressed versus all other cells). Top 20 correlated genes were highlighted as well as genes encoding known immune checkpoint molecules (LAG3 and TIGIT).

d) Representative images of multiplex immunofluorescence (IF) staining of CD8+LAG3+ T cells in ONB tumor tissue. Proteins detected using respective antibodies in the assays are indicated on top. Scale bars, 10 μm.

e) Developmental trajectory of CD4+ T cells with color gradient indicating the expression level of signatures related to Treg function (BATF, CTLA4, LAYN, TNFRSF9).

f) Scatter plot showing the specificity scores of regulons of CD8-PDCD1-Tex, CD8-ZNF683-Trm, CD8-GZMK-Tem, CD8-GZMH-Teff, CD4-IL21-Tfh, CD4-IL7R-Tcm, CD4-FOXP3-resting Treg, and CD4-CTLA4-suppressive Tregs. The top 10 regulons are highlighted.

g) Violin plots displaying expression levels of Exhaustion and Cytotoxic Scores in CD8+ T cell subclusters and Naiveness and Treg Scores in CD4+ T cell subclusters in ONB tumor microenvironment.
