## Supplementary material for "Single-cell transcriptomic landscape deciphers novel olfactory neuroblastoma subtypes and intratumoral heterogeneity": Figure S6

### Supplementary Figure S6

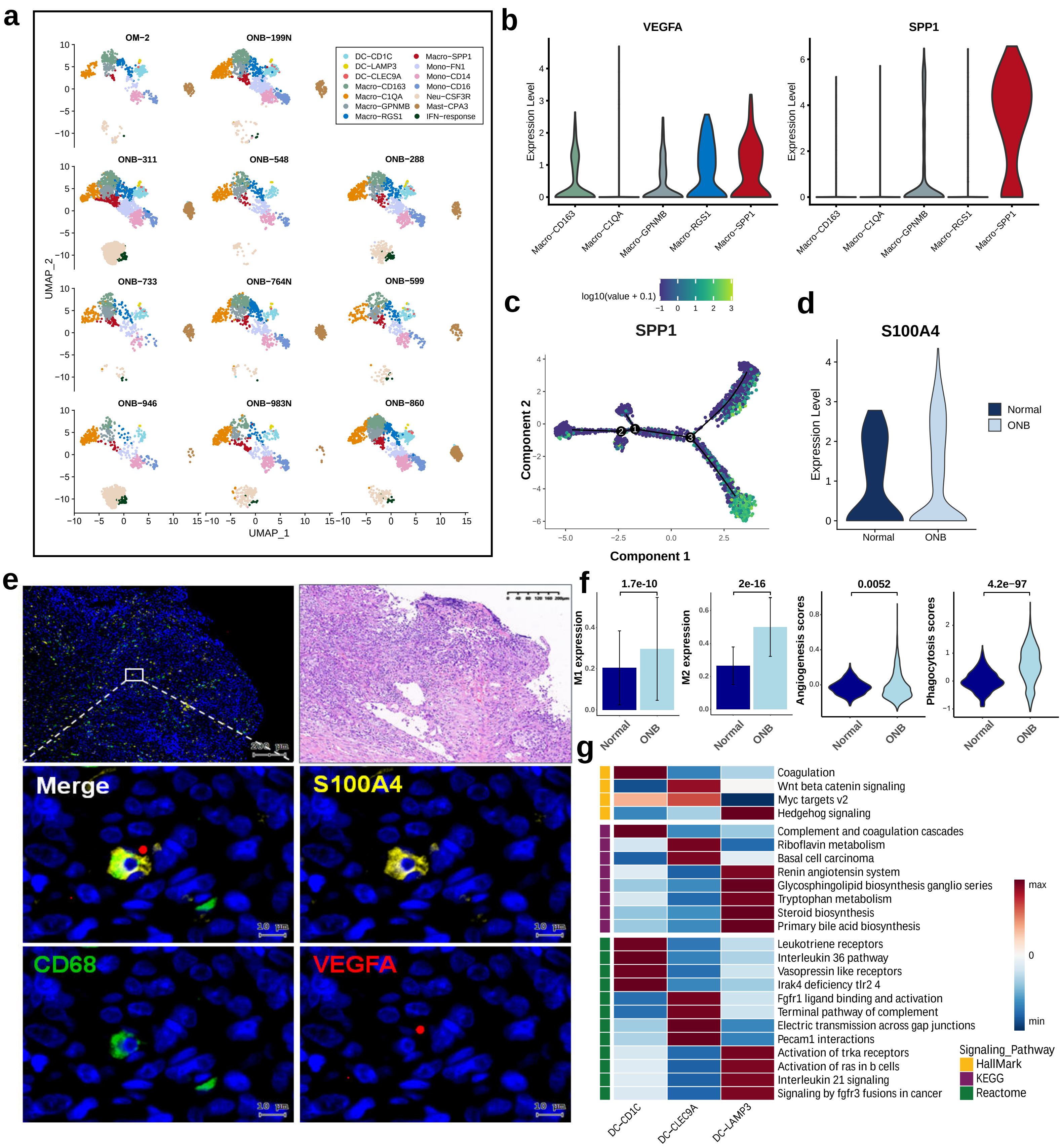

**Supplementary Figure S6.** scRNA profiles of myeloid subclusters in ONB tumor microenvironment.

a) UMAP visualization of macrophages, DCs and monocyte subclusters split by patient origins, colored by cell subtypes.

b) Violin plots displaying expression levels of marker genes in macrophage subclusters in ONB tumor microenvironment.

c) Developmental trajectory of macrophages with color gradient indicating the expression level of M2 TAM signature SPP1.

d) Violin plots showing the distribution of S100A4 in macrophages, stratified by normal and tumor tissue origins.

e) Representative images of multiplex immunofluorescence (IF) staining of macrophages in ONB tissues. Proteins detected using respective antibodies in the assays are indicated on top. Scale bars, 10 µm.

f) Boxplots and violin plots comparing M1 & M2 Scores, Phagocytosis & Angiogenesis Scores among normal and ONB derived macrophages. P values were calculated using the Wilcoxon rank-sum test.

g) Heatmap showing the selected signaling pathways that were significantly enriched in GSVA analyses for each DC subcluster derived from Hallmark, KEGG and Reactome signaling pathways. Filled colors from dark-blue to red represent scaled expression levels (normalized -log10 p values) from low to high.
