## Supplementary material for "Single-cell transcriptomic landscape deciphers novel olfactory neuroblastoma subtypes and intratumoral heterogeneity": Figure S7

### Supplementary Figure S7

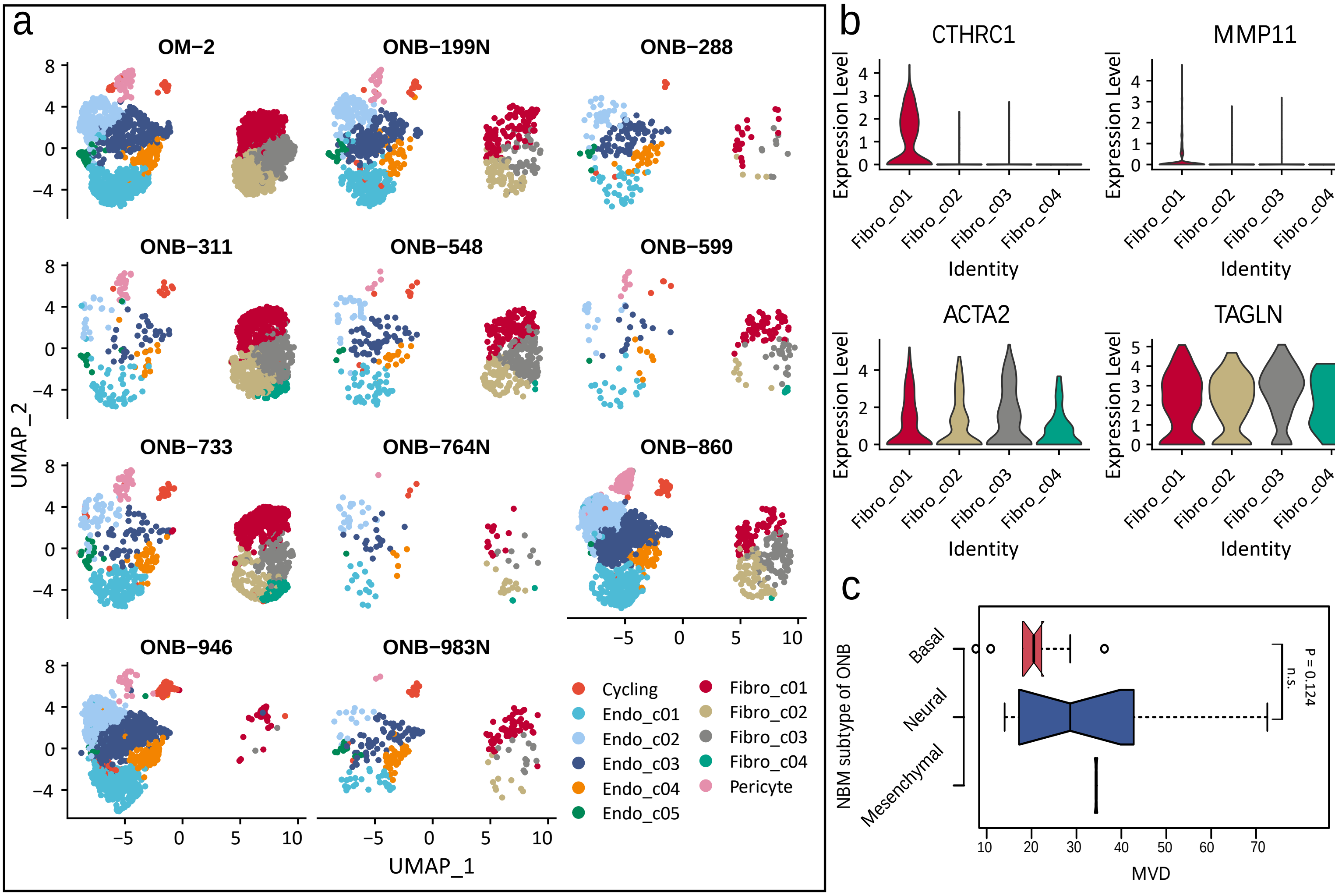


Supplementary Figure S7. scRNA profiles of stromal subclusters in ONB tumor microenvironment.

a) UMAP visualization of EC and fibroblast subclusters split by patient origins, colored by cell types.

b) Violin plots displaying expression levels of marker genes in fibroblast subclusters in ONB tumor microenvironment.

c) Boxplots comparing MVD levels among Basal, Neural and Mesenchymal ONB subtypes. P value was calculated using the one-way ANOVA test. n.s., no significance.
